# Native architecture, allosteric modulation and gating mechanism of glycine-dependent NMDA receptors

**DOI:** 10.64898/2026.04.13.718261

**Authors:** Junhoe Kim, Avery J. Benton, James S. Lotti, Nirvan Rouzbeh, Kasper B. Hansen, Eric Gouaux

## Abstract

*N*-methyl-D-aspartate receptors (NMDARs) mediate excitatory signaling essential for synaptic plasticity and memory. Unlike GluN2-containing NMDARs, GluN3-containing receptors are activated solely by glycine, exhibit profound desensitization and paradoxical potentiation by GluN1-selective antagonists, including CGP-78608 (CGP). Although GluN3 NMDARs regulate synapse pruning, excitotoxicity, and are associated with schizophrenia, autism and stroke, their native stoichiometry and gating mechanism poorly defined. Using single-molecule pulldown analysis, we show that native GluN3A receptors are diheteromeric assemblies. Cryo-EM analysis of GluN1/GluN3A receptors in antagonist-bound, pre-active, active, and desensitized states, augmented by electrophysiology and pharmacology experiments, show how glycine activates the receptor solely via GluN3A-dependent conformational changes, opening the gate with ∼2-fold symmetry, and induces a ∼4-fold symmetric desensitized state. CGP-bound GluN1 restricts GluN3A rotation, promoting glycine-induced activation by blocking desensitization. These findings illuminate how CGP potentiates GluN3A receptor activity, place the receptor gating mechanism on a solid structural foundation, and define the molecular basis for pharmacological modulation.

## Main Text

NMDARs, members of the ionotropic glutamate receptor (iGluRs) family, are ligand-gated ion channels that mediate excitatory neurotransmission and are fundamental to the development and function of the nervous system^1^. NMDARs assemble as functional heterotetramers, comprising two obligatory glycine-binding GluN1 subunits and two glutamate-binding GluN2A-D and/or glycine-binding GluN3A-B subunits, and are organized into layers known as the amino-terminal domain (ATD), ligand-binding domain (LBD), and transmembrane domain (TMD)^2,3^. Activation of canonical GluN1/GluN2 NMDARs requires binding of both glycine and glutamate, resulting in channels with high Ca^2+^ permeability^1^. In contrast, GluN1/GluN3 receptors do not bind glutamate, exhibit low Ca^2+^ permeability and reduced sensitivity to pore blockers such as Mg^2+^, memantine, and MK-801^4–6^. Furthermore, agonist binding to GluN1/GluN3 receptors triggers rapid and profound desensitization^7–10^. Intriguingly, several GluN1-selective competitive antagonists potentiate GluN1/GluN3 currents by preventing glycine binding to GluN1 subunits and attenuating desensitization^11^, with CGP-78608 (CGP) delivering the most pronounced effect^12,13^. However, the structural basis underlying this paradoxical potentiation mechanism remains largely unknown.

GluN3A subunits show tightly regulated spatiotemporal expression, with elevated levels during early postnatal development that promote synaptic pruning and maturation^14–17^. In adults, GluN3A persists in limbic structures involved in emotional regulation and stress responses^18,19^. Dysregulated GluN3A expression or function has been implicated in schizophrenia, autism, addiction, and depression^4,20–22^. By contrast, GluN3B expression gradually increases during development, albeit remaining low and restricted to specific brain regions such as motoneurons^5^. Interestingly, while triheteromeric GluN1/GluN2/GluN3A receptors were initially proposed as the native assembly^4,23,24^, accumulating evidence suggests that diheteromeric GluN1/GluN3A assemblies also exist in the brain^12,25–27^. Currently, the actual subunit composition and stoichiometry of native GluN3A-containing receptors have not been definitively determined.

To date, structural and mechanistic understanding of NMDARs has focused predominantly on “canonical” GluN1/GluN2 receptors^28–37^. Agonist binding induces closure of the bilobed LBD ‘clamshell’^28–30^, producing tension through the D2–M3 linkers to open the channel gate^29,31,32,34^. These activation mechanisms are conserved across GluN1/GluN2, AMPA, and kainate receptors. During desensitization, unlike AMPA and kainate receptors that exhibit rupture of the D1–D1 LBD dimer interface^38–40^, the GluN1/GluN2 NMDARs maintain an approximate 2-fold symmetry in the ‘local’ LBD dimer under the continuous presence of agonist^29,33^. However, these canonical models do not explain how in the presence of GluN1 antagonists, binding of glycine to GluN3A subunits alone drives channel opening in GluN1/GluN3A receptors. Furthermore, the mechanism underlying profound desensitization and very slow recovery of GluN3A-containing receptors remain poorly understood.

Despite essential physiological roles and distinctive biophysical and pharmacological properties, structural and mechanistic understandings of GluN1/GluN3A receptors are unresolved, in part due to the absence of high-resolution structures across the receptor gating landscape. A recent study reported two engineered GluN1/GluN3A structures, bound to the non-selective antagonist CNQX or to glycine, both resolved at medium resolution^41^. These constructs incorporated cysteine substitutions within the GluN1 M4 and GluN3A M1 transmembrane helices for crosslinking, yet densities for these regions were unresolved, and density for the selectivity filter structure was also absent, resulting in an incompletely defined ion channel pore (Supplementary Fig. 1). Moreover, because CNQX binds to GluN1 and GluN3A LBDs with similar affinities^42^, mechanistic conclusions from these closed inactive states explain neither glycine-driven channel opening nor do they address the crucial question of CGP potentiation. Here, we determine the subunit stoichiometry of GluN3A-containing NMDA receptors from the native mouse brain by single-molecule pulldown analysis using subunit-specific antibodies. Furthermore, we present multiple GluN1/GluN3A receptor structures along their gating trajectory, including pre-active and active states in complex with CGP, glycine, and positive allosteric modulators (PAMs), providing a comprehensive mechanistic scheme for GluN1/GluN3A activation and desensitization, and for receptor potentiation by CGP and modulation by PAMs.

### Native receptors harbor two GluN3A subunits

To enable isolation of native GluN3A-containing NMDARs^14,15^, we developed a high affinity, highly specific GluN3A monoclonal antibody, denoted 5E3 (Extended Data Fig. 1)^43^. GluN3A-containing NMDA receptors, isolated via the 5E3 Fab fused to green fluorescent protein (5E3 Fab-GFP) elute at a volume consistent with a tetrameric assembly (Fig. 1a,b), as judged by fluorescence-detection size-exclusion chromatography (FSEC)^44^. Supplementing the receptor complex with an anti-GluN1 Fab (5F11)^43^ further shifts the elution volume, indicating that the GluN3A-containing assembly in addition harbors GluN1 subunits. By contrast, the anti-GluN2A mAb (3D2)^43^ does not alter the elution volume, showing that the isolated assembly does not contain detectable GluN2A subunits.

**Fig. 1:**
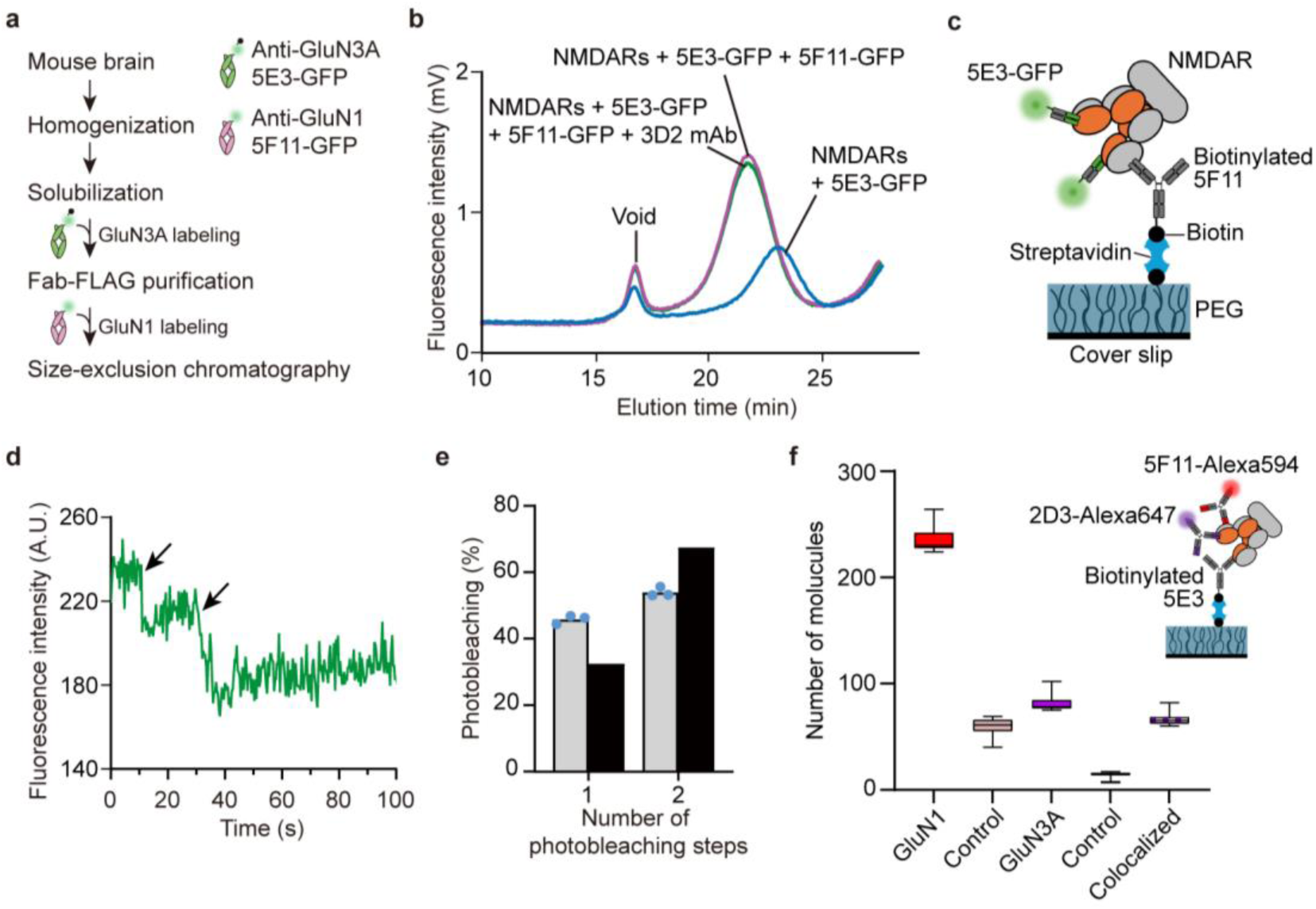
Native NMDA receptors harbor two GluN3A subunits. **a,** Flowchart illustrating the isolation procedure of native GluN3A-containing receptors from mouse brain. **b,** FSEC profiles of native GluN3A-containing receptors and their antibody complexes. **c,** Schematic of the SiMPull assay for photobleaching analysis. GluN1 and GluN3A are colored in gray and orange, respectively. **d,** Representative trace of two-step photobleaching (arrows) of native GluN3A-containing receptors labeled with 5E3 Fab-GFP. See Extended Data Fig. 2 for original image and additional traces. **e,** Distribution of GFP photobleaching steps for native GluN3A-containing receptors (gray bars) and binominal distribution (black bars) assuming 80% GFP maturation (*n* = 1,200; 400 spots per movie, 3 movie sets). **f,** Quantification of GluN1- and GluN3A-containing assemblies, as well as colocalized GluN1–GluN3A molecules detected by SiMPull. *n* = 21 over three independent experiments. Inset: Schematic of this reciprocal SiMPull assay incorporating another GluN3A-specific antibody, 2D3.

To define the subunit stoichiometry of native GluN3A-containing receptors, we performed single-molecule pulldown (SiMPull) experiments^45^ by capturing the pool of GluN1-containing native NMDARs with the 5F11 mAb and then probing for GluN3A subunits with the 5E3 Fab-GFP (Fig. 1c and Extended Data Fig. 2a,b). Photobleaching analysis of the GFP signal revealed that approximately 67% of the fluorophores bleach in two steps, while about 33% bleach in one step (Fig. 1d,e and Extended Data Fig. 2c-e), indicating that the majority of GluN3A-containing NMDARs are composed of two GluN1 and two GluN3A subunits. In a reciprocal assay using biotinylated 5E3 mAb to capture GluN3A-containing receptors and probing with 5F11 and another GluN3A-specific antibody, 2D3^43^, ∼81% of GluN3A signals colocalized with GluN1 signals (Fig. 1f and Extended Data Fig. 2f), confirming the GluN1-GluN3A assembly in the captured receptor complexes. We note that the small fraction of GFP that photobleaches in a single step may be due to the GluN3A subunit assembling with other NMDAR subunits and that resolution of this issue is the subject of future studies. Although we obtain sufficient native GluN3A-containing NMDARs for FSEC and SiMPull experiments, the yield is insufficient for single-particle cryo-EM studies. Guided by the knowledge of native GluN3A receptor subunit composition, we turned to expression of the complex in mammalian cells.

### Capture of receptor assemblies along the gating trajectory

For expression of GluN1/GluN3A receptors in cell culture, we utilized a wild-type (WT) GluN1 construct lacking the unstructured CTD (GluN1_EM_)^33^ and a similarly CTD-truncated GluN3A construct (GluN3A_EM_). Whole-cell patch-clamp recordings from GluN1_EM_/GluN3A_EM_-expressing cells demonstrate robust activation upon glycine application in the presence of CGP (Fig. 2a) albeit with faster deactivation (τ_W_ 14 ± 1 ms) and greater desensitization (69 ± 3%) compared to WT receptors (τ_W_ 65 ± 8 ms, 48 ± 3%). Nevertheless, the GluN1_EM_/GluN3A_EM_ receptor maintains similar glycine potency and functional properties compared to the WT receptor (Fig. 2b and Supplementary Table 1). Importantly, GluN1_EM_/GluN3A_EM_ is robustly expressed in cell culture, is amenable to subsequent cryo-EM studies, and lacks the putative disulfide bond between the ion channel M1 and M4 helices in the engineered GluN1/GluN3A construct previously reported^41^.

**Fig. 2:**
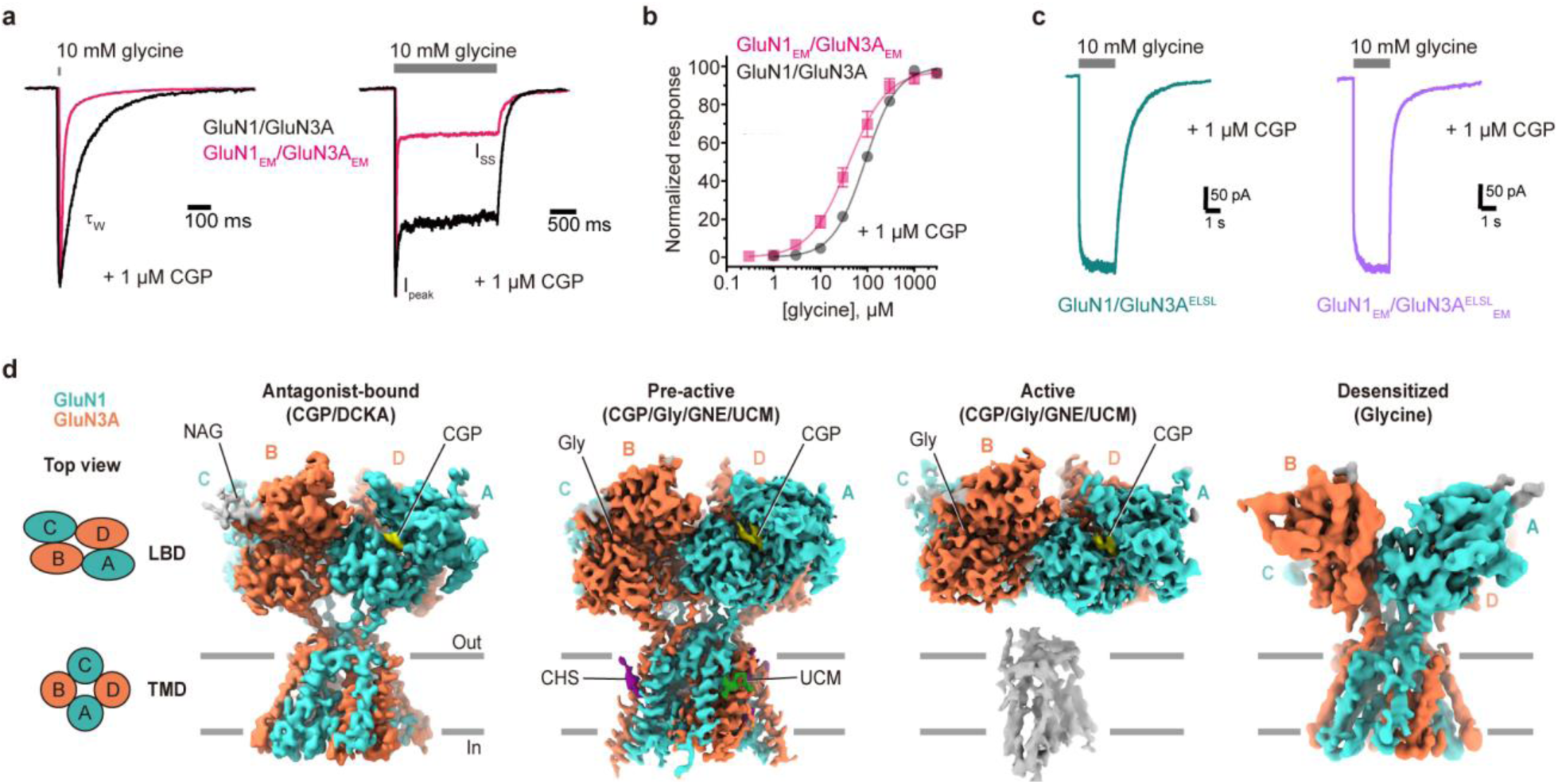
Cryo-EM snapshots capture distinct functional states. **a**, Representative whole-cell patch-clamp recordings from wild-type full-length GluN1/GluN3A and GluN1_EM_/GluN3A_EM_, normalized to peak amplitude. **b**, Glycine concentration-response data for GluN1/GluN3A (EC_50_ = 92 ± 5 µM, *n* = 9) and GluN1_EM_/GluN3A_EM_ (EC_50_ = 51 ± 9 µM, *n* = 8) in the presence of CGP. Responses were measured at steady-state using whole-cell patch-clamp recordings. Data are mean ± S.E.M. **c**, Representative whole-cell patch-clamp recordings showing attenuated desensitization in receptors containing mutant GluN3A^ELSL^ subunits. Detailed kinetic parameters and EC_50_ values are in Supplementary Table 1. **d**, Cryo-EM densities of the LBD-TMD layer of GluN1_EM_/GluN3A_EM_ receptors in the antagonist-bound (WT, CGP/DCKA), pre-active and active (ELSL, CGP/Gly/GNE/UCM), and desensitized (WT, glycine) states. Left: schematic of LBD and TMD arrangements in the antagonist-bound state, viewed from the extracellular side. Colors are: CGP (yellow), UCM (green), cholesterol hemisuccinate (CHS, purple), and *N*-acetylglucosamine (NAG, gray). The partial, low resolution TMD density in the active state is shown in gray.

To determine the structural mechanism for GluN1/GluN3A receptor gating, we sought to capture the receptor in three distinct functional states: antagonist-bound (non-active), active (open), and agonist-bound (desensitized). Because GluN1/GluN3A receptors exhibit paradoxical yet robust activation in the presence of the GluN1-selective antagonist CGP, a major aim of our work is to understand the structural mechanism of CGP potentiation. Therefore, we used CGP to occupy GluN1 subunits in both the antagonist-bound and active states. To stabilize the antagonist-bound state, we supplemented receptors with 5,7-dichlorokyurenic acid (DCKA), with the aim of blocking ambient glycine from binding to the GluN3A subunits^12,42^. For the active state, in addition to CGP and glycine, we incorporated two PAMs, GNE-9278 (GNE) and UCM-A129 (UCM), which act on GluN1 and GluN3A subunits, respectively^46,47^, and two residue mutations (described below) to further stabilize the open state of the receptor. Finally, we determined the receptor structure in a saturating concentration of glycine in the absence of CGP to capture the desensitized state.

GluN1/GluN3A receptors have a profound tendency to desensitize and multiple attempts to resolve the active, open state of the GluN1_EM_/GluN3A_EM_ receptor yielded desensitized conformations with ruptured LBD dimer interfaces. While CGP effectively slows glycine-dependent desensitization in electrophysiology experiments, it was insufficient to prevent desensitization in the cryo-EM studies. We thus stabilized the LBD dimer interface by changing GluN3A E889 and S892 to leucine, the equivalent residues in GluN2 subunits that form a hydrophobic patch^48^. The resulting GluN3A^ELSL^-containing receptor shows robust, non-desensitizing glycine currents in the presence of CGP (Fig. 2c). Moreover, GluN1/GluN3A^ELSL^ receptors exhibit increased glycine potency compared to WT receptors, suggestive of energetically favorable access to the active state (Supplementary Table 1). We therefore utilized GluN1_EM_/GluN3A^ELSL^_EM_ receptors to determine an activated, open channel conformation.

By combining pharmacological tools with the judiciously chosen mutant receptor, we resolved cryo-EM structures of GluN1/GluN3A receptors at 2.8-4.2 Å resolution in the antagonist-bound (CGP/DCKA), pre-active and active (CGP/Gly/GNE/UCM), and desensitized (glycine) states, focusing on the structural regions that govern receptor function (Fig. 2d and Supplementary Table 2). Under the CGP/Gly/GNE/UCM condition, we identified two major conformational classes that we deem pre-active and active, primarily distinguished by their TMD conformations.

All four structures displayed a ‘fuzzy’ ATD layer, indicative of conformational heterogeneity. Subsequent 3D classification and refinement focusing on the ATD layer only provided subclasses with poorly defined ATD features, consistent with the previously reported GluN1/GluN3A structures^41^. Implementation of neural-network-based analysis using cryoDRGN (see Methods for details) for the antagonist-bound (Supplementary Video 1), active (Supplementary Video 2), and desensitized (Supplementary Video 3) states revealed that the ATD conformations are continuous, limiting reliable structural modeling of ATD dimers. One implication of these findings is that the canonical allosteric modulation mediated by stable ATD-LBD interactions in GluN1/GluN2 NMDARs^49^ is absent in GluN1/GluN3A receptors.

### Antagonist-bound LBD reveals weak dimer interface

The antagonist-bound GluN1_EM_/GluN3A_EM_ receptor reveals that GluN1 and GluN3A subunits occupy the A/C and B/D positions, respectively, identified by structural features of loop1 on GluN1 and helix A’ on GluN3A (Extended Data Fig. 3)^50^. This arrangement resembles that of GluN1/GluN2 receptors, where GluN1 subunits occupy A/C and GluN2 subunits B/D positions^1^. Through focused refinement, we resolved the LBD-TMD layer of the antagonist-bound GluN1_EM_/GluN3A_EM_ receptor at ∼3.67-Å resolution, enabling fitting of LBD side chains, identification of bound antagonists, and building a complete well resolved TMD (Extended Data Fig. 4a). CGP stabilizes the GluN1 ‘clamshell’ in an ‘open’ conformation, revealing a cleft that is 22° more open than the glycine-bound LBD (PDB 2A5T, Extended Data Fig. 4b-d)^48^. In the GluN3A clamshells, however, no discernable density was observed for DCKA, despite its inclusion at 5 mM concentration (see Methods) and the clamshells adopt a conformation similar to the previously described apo conformation (Extended Data Fig. 4e)^51^.

The D1-D1 interface between receptor subunits at a ‘local’ LBD dimer plays a central role in glutamate receptor function, with an ‘intact’ interface associated with resting and active states and a ‘disrupted’ interface associated with desensitized states^1^. In the antagonist-bound state, we found that the GluN1-GluN3A D1-D1 interface is less extensive compared to that of the non-desensitized GluN1/GluN2A receptor, burying 779 Å^2^ of surface contact area versus 941 Å^2^ (PDB 2A5T)^48^, primarily due to fewer interactions between helix D on GluN1 and helix J on GluN3A, where E889 and S892 in GluN3A are positioned without compensatory interactions (Extended Data Fig. 4f-h). We speculate that in GluN3 receptors the acidic E889 and polar S892 residues destabilize the D1-D1 LBD dimer interface and that due to weaker D1-D1 interactions GluN1/GluN3A receptors undergo rapid and sustained desensitization when the LBD dimers uncouple.

### Stabilized LBD dimer primes channel opening

We utilized the GluN3^ELSL^ variant to study activated states, where the D1-D1 interface is stabilized, thereby increasing the likelihood of capturing non-desensitized states of the receptor along its activation pathway. Single-particle cryo-EM on the GluN1_EM_/GluN3A^ELSL^_EM_ LBD-TMD layer in complex with CGP/Gly and the PAMs GNE and UCM identified two major conformations: one class (∼12%; class 1) shows a well resolved LBD and TMD, while a second class (∼88%; class 2) exhibits a well resolved LBD but diffuse density for the TMD, limiting structural modeling (Fig. 3a). We proceeded to refine class 1 at 3.01-Å resolution, allowing identification of both CGP and glycine, key side chain densities, and intact LBD-TMD linkers (Extended Data Fig. 5). The well-resolved TMD structure of class 1 defines the gating ring formed by GluN1 V656 and GluN3A V769 (V-gate), equivalent to the VIVI-gate in GluN2-containing NMDARs^52^. The V-gate in class 1 is wider at the B/D positions relative to the antagonist-bound state, although the pore remains closed.

**Fig. 3:**
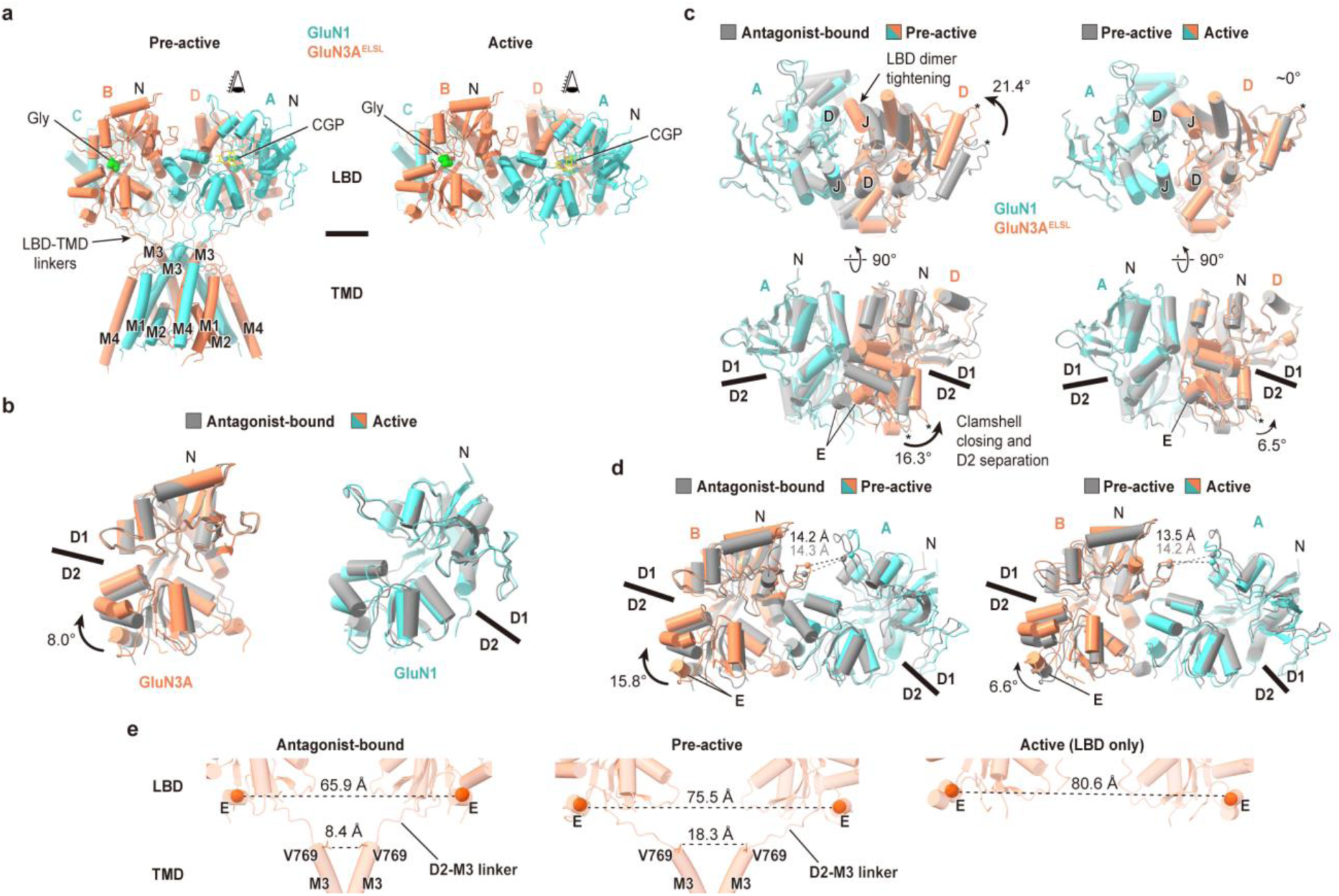
Stabilized LBD dimer primes channel opening. **a**, Side views of the pre-active (class 1) and active (class 2) GluN1_EM_/GluN3A^ELSLEM^ structures. CGP (yellow) and glycine (green) are shown as stick and sphere models. **b**, Superposition of GluN3A^ELSL^ and GluN1 LBDs from antagonist-bound (gray) and active (colored) states, aligned by D1 lobe Cα atoms. **c**, Superposition of GluN1-GluN3A^ELSL^ LBD dimers between antagonist-bound and pre-active (left), and pre-active and active (right) states, aligned at GluN1 D1. Views shown from the top (‘eyes’ in **a**) and side. Rotations of the GluN3A^ELSL^ LBD were measured as angles between COMs of S564 (asterisks), referenced to the LBD COM. D2 separations were similarly measured using P836. **d**, Comparison of LBD interdimer structures between antagonist-bound and pre-active (left), and pre-active and active (right) states, aligned at the LBD layer. Rotations were measured as angles between COMs of the D2 lobes and the LBD in GluN3A^ELSL^. Distances were measured between COMs of GluN1 R489 and GluN3A E534. **e**, LBD-TMD conformations of GluN3A in the antagonist-bound, pre-active, and active states, with distances measured between COMs of helix E or the closest carbon atoms of V769.

In class 2, the LBD layer was resolved at 2.81-Å resolution with local resolution reaching ∼2.3 Å, revealing glycine interactions in the GluN3A^ELSL^ LBD cleft, consistent with the isolated GluN3A LBD crystal structure (Extended Data Fig. 6a)^50^. CGP was resolved in the GluN1 LBD cleft in an unambiguous orientation (Extended Data Fig. 6b). Despite the differences in the TMD resolutions, the GluN3A clamshells are nearly identical in both classes, showing ∼8° more closed conformations compared to the apo state (Extended Data Fig. 6c and Fig. 3b). Based on their clamshell structures and ligand occupancies, we conclude that the LBDs in class 1 and 2 are associated with ‘activated’ states of the receptor. We suggest that class 1, where we can visualize the TMD and ion channel gate, represents a constrained, pre-active conformation and that class 2 corresponds to an active conformation.

Further image processing of single particle cryo-EM data collected on receptor complexes in the presence of CGP/Gly/GNE/UCM uncovered another 3D class, deemed class 3, in which portions of the TMD are resolved. Notably, class 2 and class 3 adopt indistinguishable LBD conformations. Although the local resolution is low, and the density is not sufficient for building an atomic model, the location and features of the additional TMD density in class 3 enable th identification of the extracellular portions of the M3 helices from all subunits, the region of the TMD that includes the ion channel gate. (Extended Data Fig. 6d-f). Comparison of the density features of class 3 with the pre-active state (class 1) shows a pronounced displacement of the B/D M3 helices away from the central axis and subtle movement of the M3 helices for the A/C subunits, mirroring the conformational changes within the LBD layer, where the B/D subunits under large movements and the A/C subunits only small displacements. Taken together, we suggest that class 3 corresponds to an open conformation, in which the cross section of the pore, near the extracellular side of the membrane, is a 2-fold symmetric parallelogram and thus distinct in shape and symmetry from the open gate cross sections of GluN1/GluN2 NMDARs, as well as AMPA and kainate receptors. We also speculate that the narrower cross-section of the GluN1/GluN3A receptor pore in the A/C direction, together with the positively charged arginine lining the GluN3A pore, could explain its low channel conductance^25^.

Activation of GluN1/GluN2, AMPA, and kainate receptors is achieved by conformational rearrangements in the LBD layer that require a stable LBD D1–D1 interface to couple the agonist-induced clamshell closure and rotation of the LBD dimers to the opening of the ion channel gate^1,34^. Here we harness the GluN1-GluN3A^ELSL^ variant to stabilize the D1-D1 interfaces in the pre-active (class 1) and active (class 2) states, mirroring the corresponding interface in the GluN1/GluN2A structure^48^. Upon comparing the pre-active state of the GluN1-GluN3A^ELSL^ complex with the wild-type-like antagonist-bound state we find that the GluN3A^ELSL^ LBD has undergone a 21.4° rotation toward the GluN1 LBD (Fig. 3c). This conformational difference, together with glycine-induced clamshell closure, combines to reorient the GluN3A^ELSL^ D2 lobes by a ∼16.3° rotation away from the dimer interface, leading to separation of the D2–D2 domains (Fig. 3c). In the active state, while the D1–D1 interface remains unchanged from the pre-active state, the GluN3A^ELSL^ D2 lobes rotate an additional ∼6.5°, further separating the D2–D2 lobes. These movements collectively result in rotation of the GluN3A^ELSL^ D2 lobes away from each other within the LBD layer (Fig. 3d). By contrast, the relative position of the GluN1 LBD remains largely unchanged in the pre-active and active states, as shown by only subtle distance changes between GluN1 R489 and GluN3A E534 (Fig. 3d).

In GluN1/GluN2, AMPA, and kainate receptors, separation of helix E within the D2–D2 lobes transmits ‘tension’ through the D2–M3 linkers and exerts a ‘pulling’ force on the M3 helices^1^. This pulling force is more pronounced at the B/D positions than the A/C positions and serves as a structural metric for channel opening^1^. In GluN1/GluN3A receptors, conformational changes in the pre-active state result in a 9.6 Å increase in the distance between GluN3A helix E at the B/D positions compared to the antagonist-bound state (Fig. 3e). This separation leads to a corresponding 9.9 Å increase between the gate residues (V769), expanding the channel gate at the B/D positions and priming it for opening. Additional separation of the GluN3A D2 lobes in the active state induces an additional 5.1 Å increase between helix E in GluN3A subunits, which we speculate further separates the channel gate at the B/D positions. We conclude that a stable, intact LBD dimer interface in combination with closure of the GluN3A clamshell drives separation of the GluN3A M3 helices through the B/D positions to achieve channel opening of GluN1/GluN3A receptors, while the GluN1 clamshells remain in a CGP-bound, ‘blocked open’ conformation.

### LBDs adopt ∼4-fold symmetry upon desensitization

A hallmark of GluN1/GluN3A receptors is rapid and nearly complete desensitization followed by slow recovery, with time constants for desensitization and recovery of 34 ± 4 ms and 5.5 ± 0.7 s, respectively (Fig. 4a,b and Supplementary Table 1). To understand the molecular basis of this process, we elucidated the structure of the GluN1_EM_/GluN3A_EM_ receptor in the presence of saturating glycine (1 mM). Iterative single-particle data analysis did not detect multiple discernible conformational classes for the LBD and TMD layers (Extended Data Fig. 7). Focused classification and refinement on the LBD/TMD layers yielded a 4.21-Å resolution reconstruction, which enabled resolution of the LBDs and transmembrane helices, yet was insufficient to fit small side chains.

**Fig. 4:**
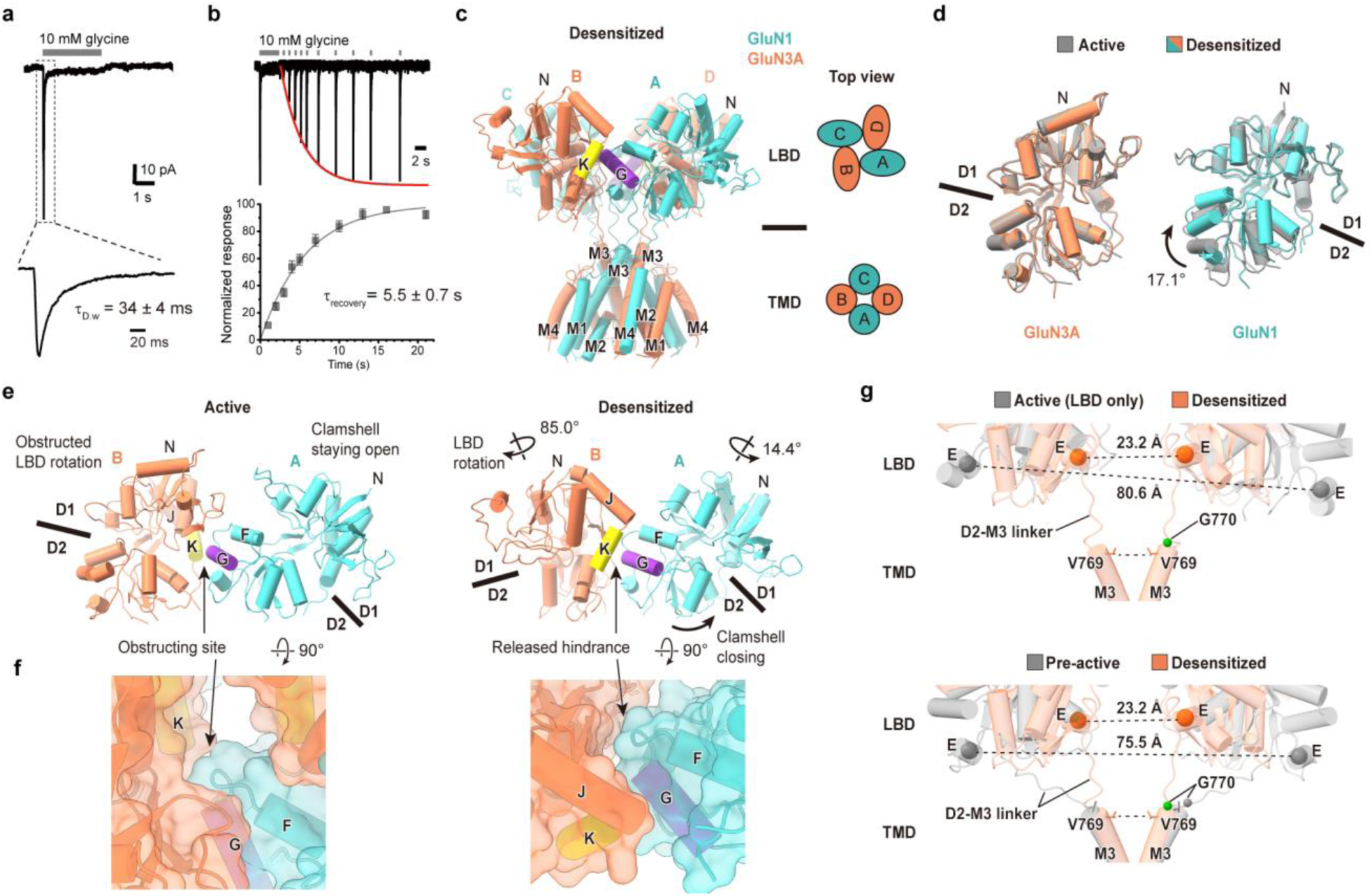
LBD rearranges upon desensitization. **a**, Representative whole-cell recording of GluN1/GluN3A receptor activated by 3 s application of glycine in the absence of CGP. Highlighted region shows desensitization time course. **b**, Overlay of recordings of GluN1/GluN3A recovery from desensitization in the absence of CGP, with increasing intervals between glycine applications. Plot summarizes normalized response amplitudes as function of time between applications (*n* = 7, mean ± S.E.M). **c**, Side view of the desensitized GluN1_EM_/GluN3A_EM_ structure with schematic of LBD and TMD arrangements in top view. **d**, Superposition of active (gray) and desensitized (colored) GluN3A and GluN1 LBDs, aligned by D1 lobe Cα atoms. **e**, Comparison of LBD layers in the active and desensitized states shows a large degree of GluN3A LBD rotation. **f**, Top view of LBD interdimer interfaces showing contacts between helix K of GluN3A (yellow) and helix G of GluN1 (blue), overlaid with surface model (transparent). **g**, Structural comparisons of the GluN3A LBD-TMD layer in the pre-active, active, and desensitized states. G770 in the D2-M3 linker, serving as a pivot point, is shown as a sphere.

The glycine-bound receptor undergoes large conformational changes from a 2-fold symmetric to a pseudo 4-fold symmetric LBD arrangement, achieved by rotation of the GluN3A LBDs (Fig. 4c). This conformation resembles the ‘deeply desensitized’ states of AMPA and kainate receptors^38–40^, but has not been observed in GluN1/GluN2 NMDARs. The dramatic reorientation of the GluN3A LBD positions helix K of GluN3A adjacent to helix G of GluN1 (Extended Data Fig. 8a), reminiscent of the kainate receptor “desensitization ring”^39,40^. In the desensitized state, the GluN1 LBD undergoes a 17.1° clamshell closure compared to its CGP-bound conformation in the active state, while the clamshell closure of the GluN3A LBD remains unchanged (Fig. 4d). Additionally, the GluN1 LBD rotates ∼14.4° ‘outward’ from the central axis, allowing the GluN3A LBD to rotate by ∼85.0° (Fig. 4e,f). At the GluN1 and GluN3A LBD interface, we identified potential salt bridges formed by GluN3A residues that are not conserved in GluN2 subunits (Extended Data Fig. 8b,c). Disruption of these interactions by mutating R704 in the GluN1 helix G, which interacts with D901 and D905 in the GluN3A helix K, to either glutamate (R704E) or alanine (R704A) accelerates recovery from desensitization (Extended Data Fig. 8d-f), consistent with the hypothesis that in the desensitized state, the receptor visits the pseudo 4-fold LBD arrangement visualized in the cryo-EM reconstruction.

The position of the GluN3A LBDs in the desensitized state reduces the distance between GluN3A helix E from the active and pre-active states (Fig. 4g), relieving tension on the M3-D2 linker, leading to channel gate closure. In comparing the pre-active and desensitized structures, we identified G770 at the ‘top’ of GluN3A M3 helix as the site of conformational changes in the D2-M3 linker upon LBD rearrangement. Mutation of G770 to glutamine, the equivalent residue in GluN2 subunits, significantly attenuates the rate of recovery from desensitization in the absence of CGP (12 s vs. 5.5 s) and nearly doubles the extent of desensitization in the presence of CGP, compared to WT receptors, respectively (Extended Data Fig. 8g-i). We suggest that G770 serves as a ‘pivot point’ in the polypeptide backbone, facilitating GluN3A LBD rotation during desensitization.

### GluN1/GluN3A channel opening is 2-fold symmetric

The closed channel gate of NMDA, AMPA, and kainate receptors is formed by a bundle of M3 helices that constrict the pore, physically occluding ion flow, and its iris-like expansion enables ion conduction through a ∼4-fold symmetric pathway^53^. A distinct feature of GluN1/GluN3A channel gating in the presence of CGP compared to GluN1/GluN2 NMDARs, AMPA, and kainate receptors is that the GluN1 clamshell remains in an open, antagonist-bound conformation throughout activation, and thus the gating is solely driven conformational changes in GluN3A subunits. Our structural analysis of GluN1/GluN3A receptors allows a direct comparison with the gating mechanism of GluN1/GluN2B receptors to understand the distinct features of GluN1/GluN3A activation (Fig. 5a-c).

**Fig. 5:**
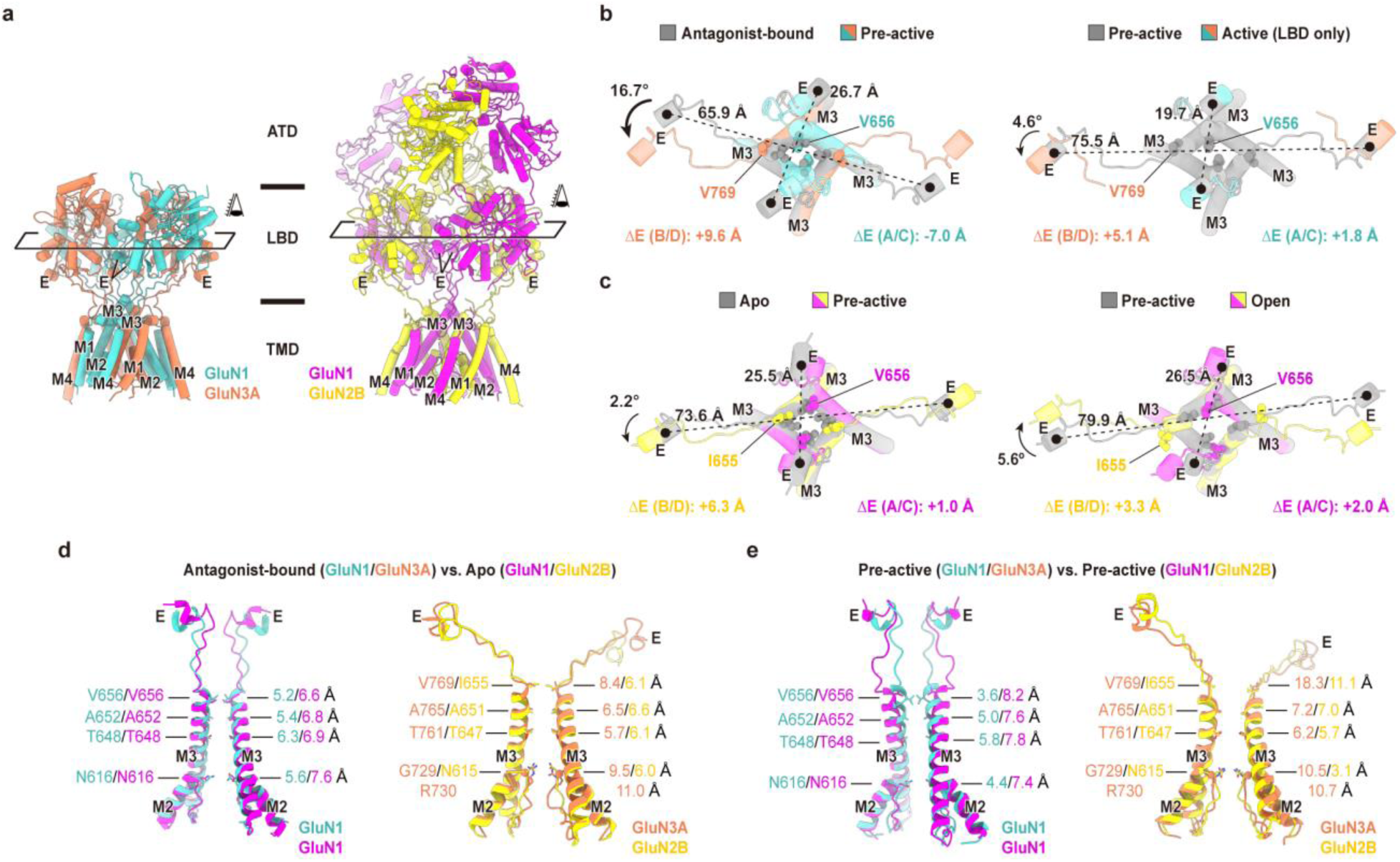
Comparing GluN1/GluN3A and GluN1/GluN2B channel opening mechanisms. **a**, Side views of the pre-active GluN1/GluN3A and open state GluN1/GluN2B (PDB 9BIB) structures. **b**, Comparison of GluN1/GluN3A channel gates between antagonist-bound and pre-active states (left), and pre-active and active states (right), viewed from top (‘eyes’ in **a)**. Gate residues are shown as spheres. COMs of helix E (black dots) were used to measure inter-subunit distances and their changes (ΔE). Curved arrows and dotted lines connecting COMs indicate relative LBD rotations between compared structures. **c**, Comparison of GluN1/GluN2B channel gates between apo (PDB 9ARG) and pre-active (PDB 6WI1) states (left), and pre-active and open states (rights), viewed from the top. **d**, Structural comparison of pore-forming helices (M2 and M3) between antagonist-bound GluN1/GluN3A and apo state GluN1/GluN2B. Distances between closest pore-lining carbon atoms are shown. **e**, Comparison of pore-forming helices between pre-active GluN1/GluN3A and pre-active GluN1/GluN2B.

To compare the CGP/Gly-bound pre-active and active structures, we utilized the Gly/Glu-bound pre-active and open GluN1/GluN2B structures^34,52^. Transitions from the antagonist-bound to pre-active states and from the pre-active to active states in GluN1/GluN3A involve greater distance changes between helix E in B/D positions (9.6 Å and 5.1 Å, respectively), compared to the corresponding changes in GluN1/GluN2B (6.3 Å and 3.3 Å). Notably, during GluN1/GluN3A activation, the distance between helix E of the GluN1 subunits, in the A/C positions, decreases due to ‘tightening’ of the LBD D1-D1 interface, in contrast to the corresponding slight increases in GluN1/GluN2B activation. These differences accompany distinct LBD-to-TMD reorientations during activation, characterized by a 4.6° counterclockwise rotation of the GluN3A LBDs parallel to the membrane plane, in contrast to a 5.6° clockwise rotation of the GluN2B LBDs (Fig. 5b,c). We also note that glycine-induced closure of the GluN3A clamshells results in allosteric movement of the GluN1 LBDs and an increase in the distance between their respective helix E elements of 1.8 Å, although the overall distance remains substantially smaller than that of the corresponding distance in the GluN1/GluN2B receptor (Fig. 5b,c). Inspection of ion channel regions reveals a more pronounced expansion at the channel gate for GluN3A subunits in the GluN1/GluN3A receptor in comparison to GluN2B subunits in the GluN1/GluN2B receptor (Fig. 5d,e), mirroring the larger conformational changes in the GluN3A LBD layer compared to the GluN2B LBD layer. By contrast, the extracellular portion of M3 and the channel gate in GluN1 subunits remain narrower in GluN1/GluN3A compared to GluN1/GluN2B receptors (Fig. 5d,e). These structural observations are consistent with previously reported results using the substituted cysteine-accessibility method that suggested a narrower extracellular vestibule in GluN1/GluN3A compared to GluN1/GluN2 receptors^54^.

Together, these findings suggest that GluN1/GluN3A channel opening, in the presence of CGP, proceeds with approximate 2-fold symmetry, in contrast to the ∼4-fold symmetry observed in GluN1/GluN2 NMDA, AMPA, and kainate receptors. We speculate that the larger separation of the D2 lobes of GluN3A subunits, by contrast with GluN2B subunits, compensates for the absence of GluN1 clamshell closure during activation of GluN1/GluN3A receptors and enables opening of the ion channel gate.

### Molecular architecture of the ion channel

The previously reported crosslinked GluN1/GluN3A TMD lacked the structure of selectivity filter and the GluN1 M4 and GluN3A M1 helices^41^. In contrast, our high-resolution pre-active structure resolves the GluN1/GluN3A TMD with unprecedented clarity, providing a nearly complete ion channel architecture, including the intact LBD–TMD linkers. The well-defined cryo-EM density enabled confident modeling of channel gating residues in both GluN1 and GluN3A (Fig. 6a). Importantly, the arginines at the N+1 site of GluN3A project into the channel vestibule, providing a structural basis for the significantly reduced Ca^2+^ permeability and Mg^2+^ block in GluN3A-containing receptors due to charge-charge repulsion (Fig. 6a). We also identified cholesterol hemisuccinate (CHS) molecules nestled within the pockets formed by GluN3A M1, M2, and M4, and neighboring GluN1 M1 (Fig. 6b), consistent with lipid and neurosteroid binding sites observed in GluN1/GluN2 receptors^55,56^.

**Fig. 6:**
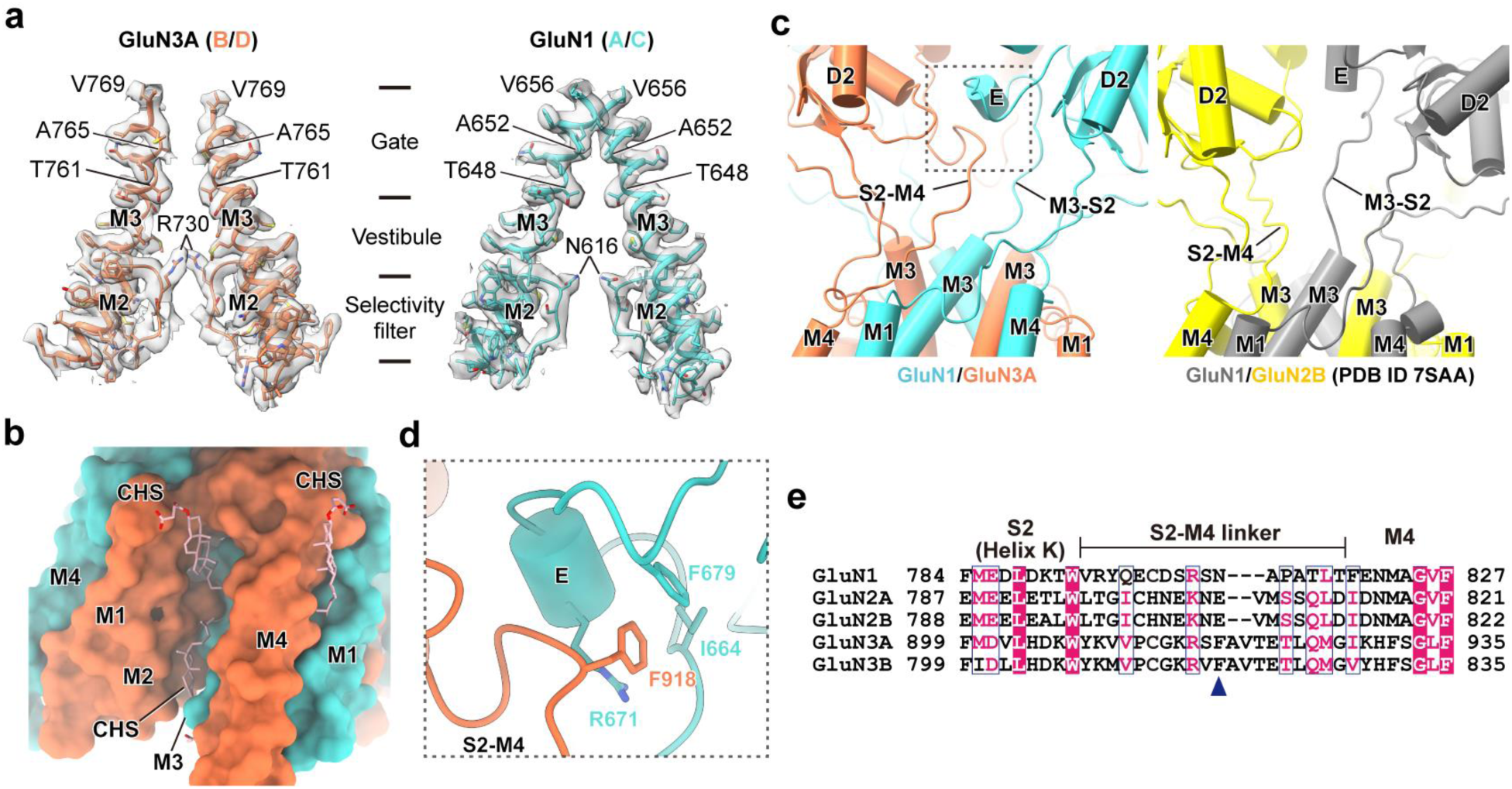
Molecular architecture of the ion channel. **a**, Cryo-EM densities (transparent) overlaid with structural models (colored) of the M2 and M3 helices in the pre-active state. **b**, Surface representation of the TMD shows CHS binding pockets formed by GluN3A M1, M2, M4 and GluN1 M1. **c**, Structures of the LBD-TMD linkers in GluN1/GluN3A (left) and GluN1/GluN2B (right) demonstrating a distinctive GluN3A S2-M4 linker conformation. **d**, The interface between the GluN3A S2–M4 linker and the GluN1 helix E (boxed region in c) shows contact residues. **e**, Sequence alignment of the S2–M4 linker across GluN1, GluN2A/B, and GluN3A/B. Blue triangle indicates the residue location of GluN3A F918.

The LBD–TMD linkers play crucial roles in NMDA receptor channel opening^1^, yet their complete structure in GluN1/GluN3A receptors has not been previously resolved. Our intact GluN1/GluN3A LBD-TMD linker structure reveals that the GluN3A S2-M4 linker adopts a conformation distinct from the equivalent region in GluN1/GluN2B (Fig. 6c). This linker forms van der Waals contacts with the helix E of the adjacent GluN1 subunit, involving GluN3A F918 and GluN1 I664, F679, and R671 (Fig. 6d). Notably, F918 is not conserved in GluN1 or GluN2 subunits, and GluN3A contains additional residues that extend the linker (Fig. 6e). These sequence differences likely contribute to the distinct GluN1/GluN3A linker architecture and may influence channel activity.

### PAMs stabilize the open state via distinct TMD interactions

PAMs exert allosteric effects by stabilizing conformations that favor channel opening^36,37,46^ or by reducing desensitization^12^. For GluN1/GluN3A channel opening, we utilized GNE and UCM, which potentiate glycine currents in both WT and mutant GluN1_EM_/GluN3A_EM_ receptors (Fig. 7a-c). In the presence of CGP and glycine, GNE and UCM potentiate GluN1_EM_/GluN3A^ELSL^_EM_ currents by ∼3.7-fold and ∼6.7-fold, respectively, and their combined application results in an ∼8.6-fold potentiation, suggestive of distinct binding sites.

**Fig. 7:**
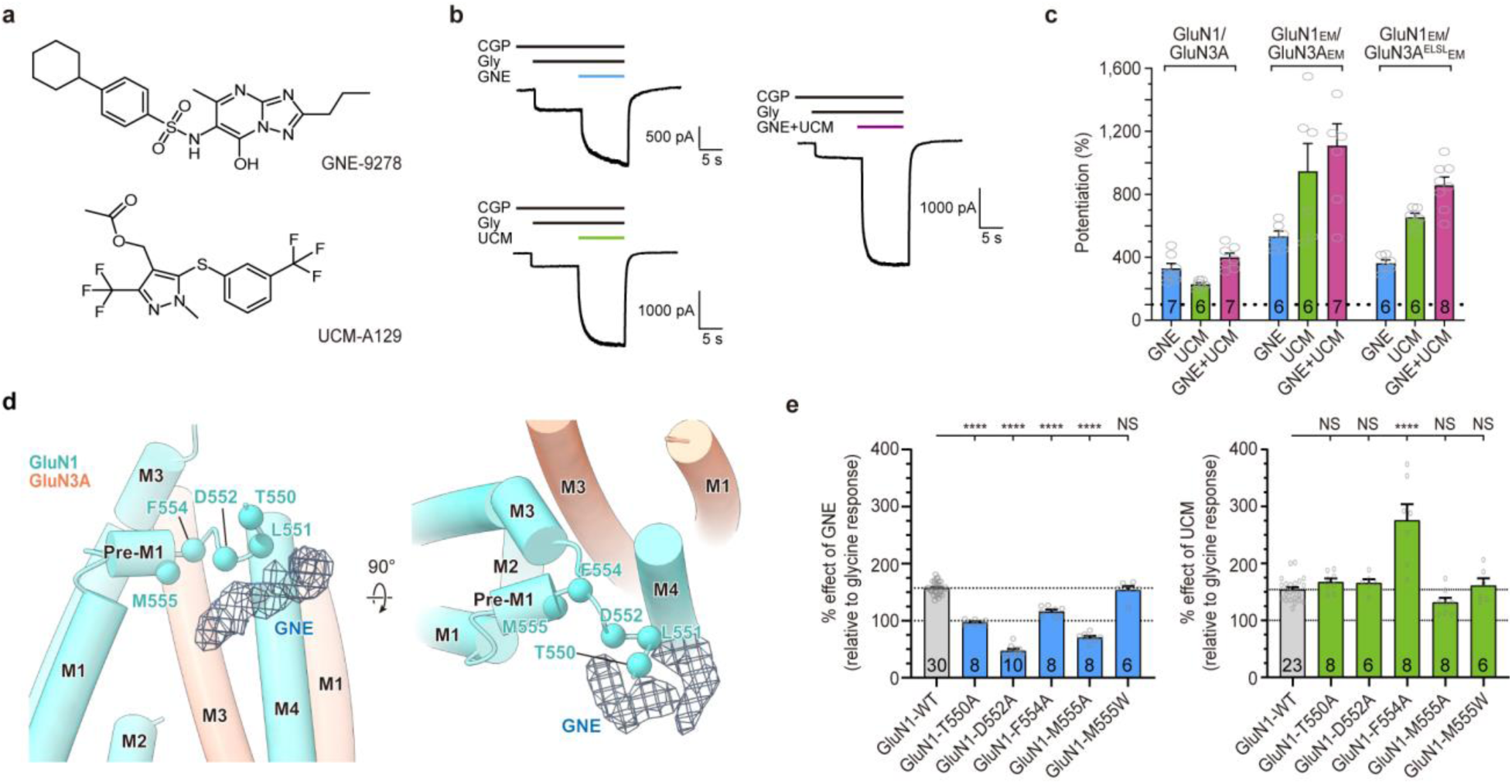
PAM binding stabilizes channel opening. **a**, Chemical structures of GNE and UCM. **b**, Representative whole-cell patch-clamp recordings from GluN1_EM_/GluN3A^ELSL^_EM_ receptors with 1 µM CGP and 1 mM glycine, in the presence of GNE (30 µM, top), UCM (100 µM, middle), or both (bottom). **c**, Bar graphs summarizing potentiation (I_PAM_/I_gly_) of WT GluN1/GluN3A, GluN1_EM_/GluN3A_EM_, and GluN1_EM_/GluN3A^ELSL^_EM_ by GNE, UCM, or their combination. *n* values are shown within bars. **d**, Partial structural model of the active-state TMD showing GNE density (blue mesh) near GluN1 pre-M1 residues (spheres). **e**, Effects of GNE and UCM on WT and mutant GluN1/GluN3A receptors measured by two-electrode voltage-clamp recordings. Statistical significance: \*\*\*\**p* < 0.0001 (one-way ANOVA with Tukey posttest).

Focused refinement on the active-state TMD layer resolves receptor TMD density with an additional, non-protein feature ‘hugging’ the GluN1 pre-M1 helix. The density feature is consistent with a bound GNE molecule and flanked by residues previously implicated in GNE binding (Extended Data Fig. 9a and Fig. 7d)^46^. Although the resolution was insufficient to define detailed molecular interactions, analysis of the protein main chain indicates that GNE binding induces conformational changes in the GluN1 pre-M1 region, shifting the linker away from the channel gate thereby creating space for M3 rearrangement (Extended Data Fig. 9b), in a fashion analogous to the PAM-induced rearrangements in GluN2B^52^. Mutations of GluN1 pre-M1 residues reveal that T550, D552, and M555 selectively impair GNE potentiation without affecting potentiation by UCM, further substantiating the molecular determinants of GNE-specific interactions (Fig. 7e). Notably, these GluN1 pre-M1 mutations do not alter glycine potency (Extended Data Fig. 9c,d and Supplementary Table 3), consistent with channel opening being primarily mediated by conformational changes in the GluN3A subunits.

Structural analysis of the pre-active state reveals a density adjacent to GluN1 pre-M4, M4, and GluN3A M1 helices, approximately the size of a UCM molecule (Extended Data Fig. 9e,f). The absence of an equivalent density in the same region in other states suggests a UCM binding site on the periphery of the TMD—a region implicated in regulating NMDAR activation efficiency^57^. Despite the predicted UCM binding site being distinct from the GNE site, the GluN1 F554A mutation reduces GNE efficacy while enhancing UCM activity (Fig. 7e), indicating that modifications at the GNE site can influence UCM modulation. Together, our findings suggest that GNE and UCM stabilize GluN1/GluN3A channel opening via distinct TMD interactions.

### CGP-bound GluN1 blocks desensitization

The resolution of GluN1_EM_/GluN3A_EM_ structures in multiple ligand-bound states provide insights into the paradoxical effects of CGP on GluN1/GluN3A receptor function (Fig. 8). In the antagonist-bound structure, where the GluN1 clamshell is stabilized in the open conformation by CGP binding, the GluN3A clamshell remains unbound and the D1-D1 interface is partially disrupted. CGP binding traps GluN1 clamshells in an ‘open’ conformation, hampering GluN3A clamshell rotation via steric hindrance between helix K on GluN3A and helix G on GluN1, thereby blocking entry of receptors into a desensitized state. Upon glycine binding to GluN3A LBDs, the clamshells stably close, resulting in the concomitant separation of helix E in D2 lobes, ‘pulling’ D2-M3 linkers and opening the ion channel gate. When glycine replaces CGP in GluN1, the GluN1 clamshells close, thereby providing sufficient space for GluN3A LBD rotation. This rotation results in receptor desensitization by releasing tension on the D2–M3 linkers and allowing channel closure. The 4-fold symmetric LBD arrangement is stabilized by interactions between LBDs and within LBDs, by bound glycine molecules. Upon glycine unbinding, the LBDs realign into their 2-fold symmetric assemblies, the D1-D1 interfaces reform, and the receptor returns to an apo, resting state.

**Fig. 8:**
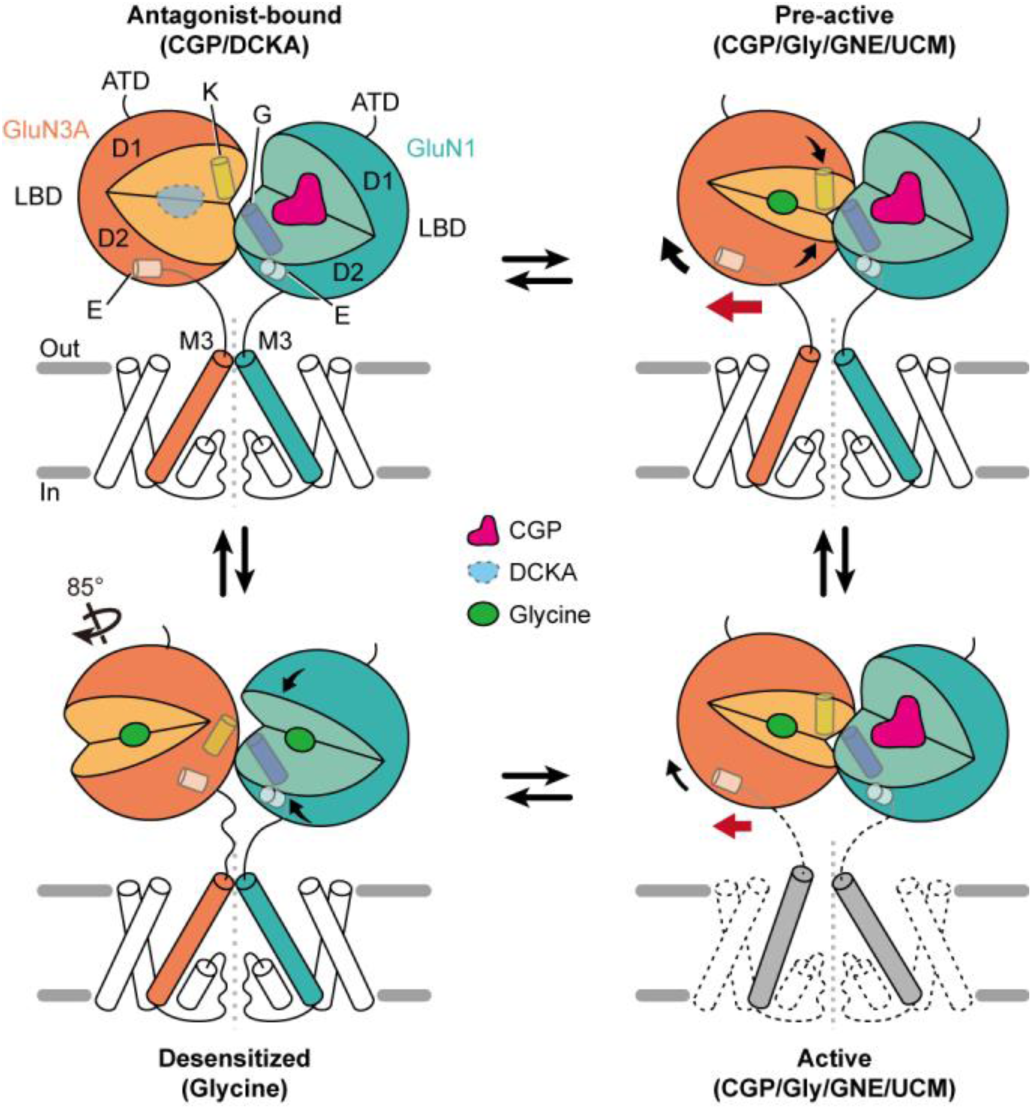
CGP modulation mechanism in GluN1/GluN3A gating. CGP binding traps the GluN1 LBD in an open conformation, stabilizing the GluN3A LBD via helix K–helix G interactions. This enables glycine-bound GluN3A LBD to efficiently transmit tension through D2-M3 linkers, priming the channel gate for opening. Replacement of CGP with glycine induces GluN1 clamshell closes, removing the structural pedestal that prevents GluN3A LBD rotation. This allows LBD reorientation into a pseudo 4-fold symmetric arrangement, relieving tension on the D2–M3 linkers and promoting channel closure. Glycine unbinding further reorients the LBDs into a 2-fold symmetric resting state, underpinning the slow recovery from desensitization. Unresolved TM elements and ligand densities are shown as dotted lines. TMD density without a structural model is in gray.

## Discussion

Here we employ a combination of fluorescence, cryo-EM and electrophysiology experiments to elaborate the native subunit composition and molecular mechanism of glycine-activated, GluN3A-containing NMDARs. Using single molecule pull down methods, we demonstrate that di-heteromeric GluN1/GluN3A receptors are the predominant physiological form in the neonatal mouse brain, a developmental stage marked by peak GluN3A expression for synaptic development and refinement^15^. Having established subunit stoichiometry of endogenous assemblies, we proceed to uncover subunit arrangement and gating mechanism of diheteromeric GluN1/GluN3A NMDARs by elucidating their conformations in key functional states and by comparing structural transitions along the gating trajectory (Supplementary Video 4). While the GluN1/GluN3A LBD organization is conserved with GluN1/GluN2 receptors, polar residues in the GluN3A LBD dimer interface disrupt a key hydrophobic patch observed in GluN2 subunits, resulting in a weak D1 dimeric interface and loss of subunit interactions during activation, thus providing a molecular explanation for the propensity of the receptor to readily desensitize. Upon desensitization, the GluN1/GluN3A LBD undergoes a pseudo 4-fold symmetric rearrangement, reminiscent of conformations observed in agonist-bound AMPA and kainate receptors^58,59^. The glycine-bound desensitized conformation is stabilized by inter-subunit interactions, consistent with the profound desensitization of GluN1/GluN3A NMDARs and the slow rate of recovery from desensitization.

A previous study has reported structures of the GluN1/GluN3A receptor in CNQX and glycine-bound states using an engineered GluN1/GluN3A construct cross-linked by non-native cysteine residues proximal to the receptor TMD^41^. These two structures were resolved at moderate resolutions (>4.1 Å)^41^, largely precluding the determination of specific interactions at subunit interfaces and limiting residue-level modelling accuracy. Moreover, the densities for the LBD-TMD linkers and the TMDs are either non-existent or weak, thus preventing modeling of the crucial peptides coupling the LBDs to the TMDs and not allowing for definition of the full TMD (Supplementary Fig. 1). While the ATD domains were included in the reported models^41^, we note that the density associated with them is weak to indiscernible, and thus the Q-scores^60^ are low or negative, suggestive of incorrect or unreliable modeling (Supplementary Fig. 1). Lastly, the incomplete density for the TMD prevented determination of the structure of the GluN1/GluN3A ion-conducting pore, which exhibits distinctive electrostatic properties due to GluN3A-containing arginine residues. By contrast, in the reconstructions reported here, we resolved the complete LBD-TMD linkers, the full TMD and ion channel pore structures, and determined pre-active and active conformations at high-resolution, providing complete structure-supported activation and desensitization mechanisms.

In our work, we provide the first structural insights into the paradoxical potentiation of GluN1/GluN3A currents by highly selective GluN1 antagonists such as CGP. While Michalski and Furukawa examined the receptor in complex with the antagonist CNQX, because CNQX binds with similar affinities to GluN1 and GluN3A subunits^42^, and does substantially potentiate GluN1/GluN3A receptor currents, the CNQX complex does not answer the question of CGP potentiation. We find that the CGP-bound LBD interface is more extensive than in the CNQX-bound state, indicating a more tightened D1 interface (Extended Data Fig. 10). Given that CGP exhibits the strongest GluN1/GluN3A potentiation among GluN1-competitive antagonists, this suggests that the degree of LBD dimer engagement correlates to the extent of potentiation, an observation further supported by our findings using the GluN1/GluN3A^ELSL^ receptor complex. Taken together, we show that CGP potentiates GluN1/GluN3A activation by preventing GluN3A LBD rotation through interdimer engagement (Fig. 8), a mechanism not observed in GluN1/GluN2 NMDARs. This suggests that GluN3A-containing NMDARs can be selectively targeted by allosteric modulators designed to influence this unique GluN3A LBD rotation. Additionally, and in the context of the membrane-embedded TMD, we find that binding of GNE and UCM bolsters GluN1/GluN3A activation, stabilizing channel opening via distinct interactions in the TMD layer. Insights gained in this study provide a structural framework for functional and pharmacological investigations of the glycine-activated GluN1/GluN3A receptor and facilitate the development of GluN3A-specific ligands and allosteric modulators with therapeutic potential in neurological and psychiatric disorders.

## Methods

### Construct design

The GluN1_EM_ construct was prepared by cloning residues 1-847 of rat GluN1-1a (Uniprot P35439) into a BacMam vector, with a thrombin cleavage site, EGFP, and an octa-histidine tag at the C-terminus^33^. The GluN3A_EM_ construct was prepared by cloning residues 1-970 of rat GluN3A (Uniprot Q9R1M7) into a BacMam vector, with EGFP and a Strep-tag II at the C-terminus. All mutations were prepared via site-directed mutagenesis.

### Protein expression and purification

Bacmid and baculovirus for wild-type GluN1_EM_/GluN3A_EM_ and mutant GluN1_EM_/GluN3A^ELSL^_EM_ receptor expression were generated using standard methods^61^. P2 viruses infected suspension HEK293T/17 cells at M.O.I 1:1. After 12 hours at 37 °C, 10 mM sodium butyrate was added, and the temperature was lowered to 30 °C. Cells were collected 48 hours post-infection, resuspended in 150 mM NaCl, 50 mM Tris-HCl (TBS, pH 8.0) with 1 mM phenylmethylsulfonyl fluoride (PMSF), 0.8 µM aprotinin, 2 µg/ml leupeptin, and 2 mM pepstatin A. Receptor complexes were extracted in 50 mM TBS (pH 8.0) with 1% (w/v) glyco-diosgenin (GDN), 1.65 mM cholesterol hemisuccinate (CHS), and protease inhibitors for 2 hours at 4 °C with stirring. To prevent binding of ambient glycine to GluN3A, a low affinity ligand, glutamate^42^, was added at 10 mM. The lysate was clarified by centrifugation at 200,000 g for 1 hour and filtered (0.22 µm cellulose). Solubilized receptors were bound to Strep-Tactin resin, washed with 10 column-volume 20 mM TBS (pH 8.0) supplemented with 0.05% (w/v) GDN (TBS-GDN), including 2 mM glutamate, and eluted with the TBS-GDN (pH 8.0) containing 10 mM D-desthiobiotin without glutamate. Receptors were concentrated with 1 mM ethylenediaminetetraacetic acid (EDTA) using 100-kDa cut-off concentrator, then purified by size-exclusion chromatography (SEC) on a Superose 6 Increase 10/300 GL column in TBS-GDN (pH 8.0) for the desensitized state or in TBS-GDN (pH 7.4) with 0.1 mM CHS and 1 µM CGP for the antagonist-bound and pre-active/active/open states. Peak fractions were pooled, concentrated to 4-5 mg/mL, and quantified at 280 nm.

### Determination of glycine contamination

Potential glycine contamination in purified GluN1_EM_/GluN3A_EM_ receptors was analyzed using liquid chromatography–tandem mass spectrometry (LC–MS/MS) with electrospray ionization in positive mode. Multiple-reaction monitoring (MRM) transitions monitored glycine quantification at m/z 76.1-to-48 for glycine and m/z 78-to-50 for deuterium-labeled glycine. Standards spiked with 0.1–5 µg/mL deuterium-labeled glycine were prepared to validate assay sensitivity. The glycine content in 1.6 mg/mL of GluN1_EM_/GluN3A_EM_ was below the lower limit of detection (0.05 µg/mL). Based on this, we estimate glycine contamination was less than 0.67 µM in 3.2 µM receptors. As this 4.8-fold difference exceeds the number of subunits in one receptor molecule (4 copies), we conclude that the majority of receptor subunits were unoccupied by glycine.

### Single-particle cryo-EM data collection and processing

The SEC purified sample was used for cryo-EM grid preparation. Quantifoil R2/1 200-mesh Au grids were glow-discharged for 1 min at 15 mA. For the antagonist-bound or desensitized state, 5 mM 5,7-dichlorokynurenic acid (DCKA) or 1 mM glycine was added immediately before vitrification. For the pre-active/active/open states, 1 mM glycine, 0.67 mM GNE-9278, 0.67 mM UCM-A129 were added immediately before vitrification. In all conditions, 0.1 mM fluorinated octyl-maltoside (FOM) was included to aid particle distribution. Vitrification was performed by applying 3 µL of protein solution on the grid, blotting for 3 sec with FEI Vitrobot Mark IV at 16 °C and 100% humidity, and plunging into liquid ethane cooled by liquid nitrogen.

The dataset for antagonist-bound state was collected on a 300 keV FEI Titan Krios with a Gatan K3 camera, BioContinuum energy filter, and Cs corrector in super-resolution mode (0.844 Å/pix), using SerialEM at a defocus range of −0.8 to −2.2 µm and a total dose of 50 e^-^/Å^2^. The dataset for pre-active (class1) and active (class 2) states was collected on a 300 kV FEI Titan Krios with a Thermo Scientific Falcon 4i camera and Selectris X energy filter (0.940 Å/pix), using SerialEM with a defocus range of −0.8 to −2.2 µm and a total dose of 60 e^-^/Å^2^. The dataset for open state (class 3) was collected on a 300 kV FEI Titan Krios with a Thermo Scientific Falcon 4i camera and Selectris X energy filter (0.743 Å/pix), using SerialEM with a defocus range of −0.8 to −2.0 µm and a total dose of 50 e^-^/Å^2^. The dataset for desensitized state was collected on a 300 keV FEI Titan Krios with a Gatan K3 camera and BioContinuum energy filter in super-resolution mode (0.829 Å/pix). Images were acquired with SerialEM at a defocus range of −1.0 to −2.4 µm and a total dose of 50 e^-^/Å^2^. Movie alignment, contrast transfer function (CTF) estimation, particle picking, 2D/3D classification, ab initio 3D reconstruction, non-uniform refinement^62^, and local refinement were performed in CryoSPARC v4^63^. Some 3D classification without image alignment were conducted in RELION-3^64^. Masks applied for signal subtraction and local refinement were generated in UCSF Chimera^65^. Map resolutions were determined at Fourier shell correlation (FSC) = 0.143 using two half maps in CryoSPARC. Data collection and model refinement statistics are summarized in Supplementary Table 2. Cryo-EM data processing workflows for the antagonist-bound, pre-active/active, and desensitized states are shown in Extended Data Figs 3, 5, and 7, respectively.

### Model building

An initial model of antagonist-bound GluN1_EM_/GluN3A_EM_ receptor was generated using AlphaFold-predicted structures of rat GluN1 and rat GluN3A^66,67^. The C-terminal regions were removed to match the expression construct, and the flexible ATD was excluded. The remaining LBD and TMD structures were segmented into D1, D2, and TMD, then independently rigid-body fitted into cryo-EM density maps using UCSF Chimera, followed by real-space refinement in Phenix^68^. The model was inspected and manually adjusted in *Coot*^69^, guided by the cryo-EM structure of CGP-bound GluN1 (PDB 7EOT)^35^ and the crystal structure of apo-state GluN3A LBD (PDB 4KCD)^51^. Bulky side chain densities in the cryo-EM map further assisted fitting. For weakly resolved M1 and M4 helices in antagonist-bound and desensitized states, unsharpened map and conserved bulky residues across NMDAR subunits initially guided helix positioning and rotation, which were then further aided by the high-resolution pre-active GluN1/GluN3A structure. Ambiguous densities, particularly at the bottoms of M1 and M4 helices and linkers between TM segments, were omitted. The independent LBD and TMD models were connected via LBD-TMD linkers to generate a complete coordinate. The final models were refined with secondary structure and Ramachandran restraints. The same approach was used to build the desensitized state model. The antagonist-bound state model was segmented and rigid-body fitted into desensitized state cryo-EM map. LBD-TMD linker were more carefully inspected and adjusted due to large GluN3A subunit rotations. Glycine density was ambiguous and thus omitted in the model. For the pre-active state GluN1_EM_/GluN3A^ELSL^_EM_ receptor, the antagonist-bound state model was used for rigid-body fitting with substituting E889 and S892 in GluN3A to leucine in *Coot*. Glycine was placed in the GluN3A^ELSL^ LBD cleft guided by the crystal structure of glycine-bound GluN3A LBD (PDB 2RC7)^50^. The active state model was derived from the pre-active state model by excluding the TMD and rigid-body fitting the LBD. Side chains were modeled in the guide of C2-symmetry maps. Refinement statistics were generated using sharpened maps with global *B*-factors in CryoSPARC, and model resolutions were determined at FSC = 0.5 in Phenix. Superpositions of structures were performed using UCSF Chimera and PyMOL v3.1 (Schrödinger, LLC). Contact areas of LBD dimer interfaces were calculated using PISA^70^. Figures were prepared using UCSF Chimera and UCSF ChimeraX^71^.

### CryoDRGN2 analysis

Neural network-based heterogeneous reconstruction method in cryoDRGN2.3^72^ was used to assess conformation heterogeneity in the ATD layer of antagonist-bound, active/open (class 3), and desensitized states. Input particle poses and CTF parameters were obtained from consensus refinement of the full receptor densities, including the ATD layer, in CryoSPARC v4, and were held fixed during CryoDRGN training. A total of 91,085 (antagonist-bound), 28,629 (active/open), and 532,539 (desensitized) particles were used as input which included some particles with diffuse ATD features that were ultimately not retained for the higher-resolution LBD-TMD reconstruction shown in Supplementary Table 2. Particle stacks were downsampled to 128 x 128 pixels, yielding pixel sizes of 3.38 Å, 2.97 Å, and 3.32 Å, respectively. An 8-dimensional latent variable model (z = 8) was trained for 50 epochs using a 1,024 x 3 encoder/decoder architecture. The output volumes were visually inspected and analyzed with cryoDRGN default settings. Conformational trajectories were generated by interpolation in the learned latent space, producing 133 (antagonist-bound, Supplementary Video 1), 117 (active/open, Supplementary Video 2), and 168 (desensitized, Supplementary Video 3) decoded volume maps. These maps were not further classified, refined, or manually interpolated. Morph movies were rendered in UCSF Chimera X.

### Isolation of native GluN3A-containing NMDA receptors from mouse brains

Native GluN3A-containing NMDARs were isolated from the brains of postnatal day 6–8 C57BL/6 mice or GluN3A KO mice (The Jackson Laboratory, Strain #:029974). Twelve brains were homogenized in ice-cold TBS (pH 8.0) with protease inhibitors, then disrupted by sonication (90 sec, 3 sec on, 9 sec off) on ice. Lysates were solubilized with 1.5% (w/v) GDN for 2 hours at 4 °C and clarified by centrifugation at 200,000 g for 1 hour. The supernatant was incubated with excess anti-GluN3A 5E3 Fab conjugated to GFP-3xFLAG tag and anti-FLAG M2 resin for 1 hour at 4 °C, loaded onto a gravity column, and washed with TBS-GDN (pH 8.0) supplemented with 1 µM CGP. Bound receptors were eluted by 3C protease digestion for 15 min on ice then concentrated for FSEC injection. FSEC performed on a Superose 6 Increase 10/300 GL column in TBS-GDN (pH 8.0) supplemented with 1 µM CGP.

### SiMPull

Coverslips and glass slides were prepared based on the previous protocol^45^. Streptavidin at 0.2 mg/mL was applied to the chamber with 50 µL and incubated for 5 minutes, followed by wash with 30 µL TBS. Biotinylated anti-GluN1 5F11 monoclonal antibody at 5 µg/mL in TBS containing 0.2 mg/mL bovine serum albumin (TBS-BSA) was applied to the chamber with 25 µL and incubated for 10 min, followed by wash with 30 µL TBS-BSA. Isolated native NMDARs diluted 1:50 in TBS-BSA containing 0.05% (w/v) GDN (TBS-BSA-GDN) was applied to the chamber with 25 µL and incubated for 20 min, followed by wash with 30 µL TBS-BSA-GDN. The probing antibodies at 2.5 µg/mL in TBS-BSA-GDN were applied to the chamber with 25 µL and incubated for 20 min, followed by wash with 30 µL TBS-BSA-GDN. The chamber was imaged using a Leica DMi8 TIRF microscope with an oil-immersion 100x objective. Images were captured using a back-illuminated EMCCD camera (Andor iXon Ultra 888) with a 133 x 133 µm imaging area and a 13 µm pixel size corresponds to 130 nm on the sample owing to the 100x objective. To estimate non-specific binding to the glass slide, the corresponding non-biotinylated antibody was applied to a separate chamber, while all other steps remained identical. This chamber was used to estimate background fluorescence. Fluorophore spots were counted by setting intensity thresholds of 3.6 for GFP-conjugated Fab and 4.0 for Alexa-labeled antibodies in ImageJ (FIJI). For single-molecule colocalization, two images were acquired from the same region of interest under different wavelengths, and molecules within a four-pixel radius were considered colocalized.

Photobleaching movies were acquired by exposing the imaging area for 120 sec. To count the number of GluN3A subunits, single-molecule fluorescence time traces of the GFP-tagged anti-GluN3A 5E3 Fab were generated using a custom Python script^73^. Each trace was manually scored as having one or two bleaching steps or was discarded if no clean bleaching steps could be identified. The resulting distribution of bleaching steps was plotted based on an estimated GFP maturation of 80%. A total of 1,200 molecules were evaluated from three independent movies. Scoring was verified by assessing decreases in GFP intensity between bleaching events. On average, the intensity drop of molecules that bleached in two steps was approximately twice that of those that bleached in a single step.

### Western blot analysis

Protein samples were mixed with loading buffer supplemented with 0.1 M dithiothreitol and incubated at 50°C for 10 minutes. The samples were then separated by sodium dodecyl sulfate polyacrylamide gel (4–20% or 15%) electrophoresis and transferred to polyvinylidene fluoride membranes. Immunoblotting was performed using primary antibodies at dilutions of 1:1,000 (anti-GluN1, Invitrogen 700685), 1:200 (anti-GluN3A, Alomone Labs AGC-030), or 1:1,000 (anti-GluA4, Millipore AB1508), followed by incubation with an IRDye® 800CW goat anti-rabbit IgG secondary antibody (Li-Cor 926-32211) at a dilution of 1:20,000. Protein bands were visualized using the Li-Cor Odyssey CLx system.

### Whole-cell patch-clamp recordings

Approximately 48 hours before recordings, HEK293T cells were plated on poly-D-lysine–coated glass coverslips (0.1 mg/ml) and cultured in Dulbecco’s modified Eagle’s medium (DMEM) supplemented with sodium pyruvate, GlutaMAX, 10% fetal bovine serum, 10 U/ml penicillin, and 10 mg/ml streptomycin (Thermo Fisher Scientific, Waltham, MA). Cells were transfected the next day via calcium phosphate precipitation^74^ with a 1:1:1 ratio of plasmid cDNAs encoding EGFP, rat GluN1-4a (GenBank U08267) and either rat wild-type GluN3A (GenBank U29873) or rat mutant GluN3A^ELSL^ containing E889L and/or S892L mutations. All other mutations were prepared in these constructs using standard site-directed mutagenesis protocols. Whole-cell patch-clamp recordings were performed ∼24 hours following transfection at room temperature (20°C) at a holding potential of –60 mV using an Axopatch 200B amplifier (Molecular Devices, San Jose, CA), filtered at 8 kHz (8-pole Bessel, Frequency Devices, Ottawa, IL) before acquisition with a Digidata 1322B with pClamp 10 (Molecular Devices). Recording electrodes were made from thin-walled borosilicate glass micropipettes (TW150F-4; World Precision Instruments, Sarasota, FL) prepared using a micropipette puller (P-1000, Sutter Instruments) to have a tip resistance of 2-3 MΩ and filled with internal solution containing 110 mM D-gluconate, 110 mM CsOH, 30 mM CsCl, 5 mM HEPES, 4 mM NaCl, 0.5 mM CaCl_2_, 2 mM MgCl_2_, 5 mM BAPTA, 2 mM NaATP, and 0.3 mM NaGTP (pH 7.35 with CsOH). The extracellular recording solution contained 150 mM NaCl, 10 mM HEPES, 3 mM KCl, and 0.5 mM CaCl_2_ (pH 7.4 with NaOH). For concentration-response experiments, a range of glycine concentrations were applied to lifted cells by gravity-driven perfusion through an array of glass pipettes positioned using a SF-77B Perfusion Fast-Step system (Warner Instruments, Hamden, CT). In experiments to determine activation, desensitization, and deactivation kinetics, fast-application of glycine to activate GluN1/GluN3A receptors was achieved using rapid solution exchange (0.6–0.8 ms for 10%–90% open-tip solution exchange times) on lifted cells with gravity-driven perfusion through a theta-glass pipette controlled by a piezoelectric translator (MXPZT-300; Siskiyou Corporation, Grants Pass, OR). CGP-78608 (CGP) and GNE-9278 (GNE) were purchased from Tocris Bioscience and UCM-A129 (UCM) was obtained from Key Organics.

### Two-electrode voltage-clamp recordings

Xenopus oocytes, purchased from Rob Weymouth (Xenopus 1), were maintained in a modified Barth’s solution as previously described^13^. The cDNAs encoding GluN1-4a and GluN3A subunits were combined at a 1:2 ratio and injected into oocytes 2-4 days before the recordings^13^. Recordings were performed at ambient temperatures with extracellular recording solution containing 90 mM NaCl, 1 mM KCl, 0.5 mM BaCl_2_, 10 µM EDTA, and 10 mM HEPES (pH 7.4). Voltage and current electrodes were filled with 3.0 M KCl and current responses were measured using a two-electrode voltage-clamp amplifier (OC-725C; Warner Instruments) at a holding potential of −40 mV, and filtered with a 20 Hz low-pass filter (Alligator Technologies) before data acquisition (PCIe-6321, National Instruments). Solutions were applied to the oocytes by gravity-driven perfusion through 8-modular valve positioners (Hamilton Company).

### Analysis of electrophysiological recordings

Concentration-response data were analyzed using GraphPad Prism (GraphPad Software, Boston, MA) by fitting response amplitudes to a range of glycine concentrations for each individual cell to the Hill equation:

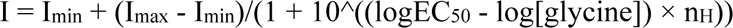

I_max_ and I_min_ are the fitted maximum and minimum current responses (I_min_ is typically negligible), EC_50_ is the agonist concentration that produces half-maximum response, [glycine] is the agonist concentration, and n_H_ is the Hill slope. For graphical representation, data points from individual cells were normalized and the averaged data points were plotted with the fitted concentration-response curve.

Recordings from fast-application whole-cel patch-clamp experiments were analyzed using Clampfit (Molecular Devices) and Axograph (https://axograph.com/). Rise times were defined by the duration from 10% to 90% of the maximal response amplitude. For recordings in the continuous presence of CGP, the desensitization time course could be adequately described using a mono-exponential fit, whereas the deactivation time course was determined using two-exponential fits to obtain τ_fast_, τ_slow_, and % fast is the fitted percentage of the fast component. For recordings in the absence of CGP, both the desensitization and deactivation time constants were determined using two-exponential fits. Weighted time constants were calculated as:

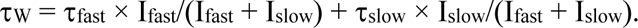

I_slow_ and I_fast_ are fitted response amplitudes for the slow and fast components, respectively.

Statistical comparisons were performed as described in figure legends using GraphPad Prism (GraphPad Software). Statistical comparisons of time constants (τ values) were performed on normally distributed rate constants (1/τ)^75^. Electrophysiological data were repeated on at least two different days as independent experiments using new recording solutions and separate transfections of the cells.

### Animal use statement

For SiMPull, photobleaching, and native receptor purification, 12 postnatal day 6-8 C57BL/6 offspring (both male and female) from Charles River Laboratories were used. Native receptor quantity per mouse was estimated via fluorescence from recombinant antibody fragments to determine the minimum required sample size. Mice were housed at 20–23°C, 40–60% humidity, with a 12:12-hour dark/light cycle. No randomization, blinding, or experimental manipulations were performed. All mice were euthanized under the OHSU Institutional Animal Care and Use Committee (IACUC) protocols, consistent with the recommendations of the Panel on Euthanasia of the American Veterinary Medical Association (AVMA) and carried out only by members of the E.G. laboratory approved on IACUC protocol TR03_IP00000905.

### Cell line statement

Sf9 cells for baculovirus production and recombinant antibody expression were obtained from Thermo Fisher (12659017, lot 421973). HEK293T/17 (tsA201) cells for protein expression and HEK293T cells for electrophysiology were from previously studies^13,33^. Cells authentication was not performed for these studies. HEK293T/17 cells tested negative for Mycoplasma contamination using the CELLshipper Mycoplasma Detection Kit M-100 (Bionique), while HEK293T cells were not tested.

## Data and code availability

Cryo-EM maps and coordinates have been deposited in the Electron Microscopy Data Bank (EMDB) and in the Protein Data Bank (PDB) under accession codes EMD-70399 and 6OEH (antagonist-bound LBD-TMD), EMD-70433 and 9OFI (antagonist-bound LBD), EMD-70446 and 9OFY (antagonist-bound TMD), EMD-70402 and 9OEM (pre-active; class 1), EMD-70403 and 9OEN (active; class 2), EMD-70404 (open; class 3), EMD-70405 and 9OEO (desensitized LBD-TMD), EMD-70437 and 9OFM (desensitized LBD), and EMD-70447 and 9OFZ (desensitized TMD), respectively. Custom code used for analyzing single-molecule photobleaching trajectories in this study is available at Zenodo (https://doi.org/10.5281/zenodo.8161179).

## Acknowledgements

We thank the Pacific Northwest Center for Cryo-EM (PNCC) supported by NIH grant R24GM154185 with assistance from Janette Myers, Cryo-EM facility at Janelia Research Campus, and OHSU Multiscale Microscopy Core (MMC) for support with microscopy, R. Courtney for assistance with manuscript preparation. We thank Atomwise Inc. for performing virtual screening and providing top hit compounds through their academic collaboration program, Artificial Intelligence Molecular Screen (AIMS). The study was supported by NIH-NINDS (R01NS116055 and R01NS097536) and NIH-NIGMS (P30GM140963) to K.B.H, NIH-NINDS (R01NS038631) to E.G, and NRF of Korea (RS202400334731) to J.K. Jennifer and Bernard LaCroute are thanked for their generous support. E.G. is an investigator of the Howard Hughes Medical Institute.

## Author Contributions

J.K. and E.G. designed the project. J.K. performed protein purification, biochemistry, SiMPull experiments, and data collection, analysis, and model building for single-particle cryo-EM. J.S.L., A.J.B, N.R., and K.B.H. designed and performed electrophysiological experiments. E.G., J.K., and K.B.H. wrote the manuscript with input from J.S.L., A.J.B., and N.R.

## Additional Information

Extended Data Figs. 1-10

Supplementary Fig. 1

Supplementary Tables 1-3

Supplementary Videos 1-4

Correspondence and requests for materials should be addressed to Eric Gouaux, or Kasper B. Hansen.

## References

1. Hansen, K.B., et al. Structure, Function, and Pharmacology of Glutamate Receptor Ion Channels. Pharmacol Rev 73, 298–487 (2021).

2. Zhou, C. & Tajima, N. Structural insights into NMDA receptor pharmacology. Biochem Soc Trans 51, 1713–1731 (2023).

3. Hanson, J.E. et al. Therapeutic potential of N-methyl-D-aspartate receptor modulators in psychiatry. Neuropsychopharmacology 49, 51–66 (2024).

4. Crawley, O., Conde-Dusman, M.J. & Perez-Otano, I. GluN3A NMDA receptor subunits: more enigmatic than ever? J Physiol 600, 261–276 (2022).

5. Chatterton, J.E. et al. Excitatory glycine receptors containing the NR3 family of NMDA receptor subunits. Nature 415, 793–8 (2002).

6. McClymont, D.W., Harris, J. & Mellor, I.R. Open-channel blockade is less effective on GluN3B than GluN3A subunit-containing NMDA receptors. Eur J Pharmacol 686, 22–31 (2012).

7. Cummings, K.A., Belin, S. & Popescu, G.K. Residues in the GluN1 C-terminal domain control kinetics and pharmacology of GluN1/GluN3A N-methyl-d-aspartate receptors. Neuropharmacology 119, 40–47 (2017).

8. Smothers, C.T. & Woodward, J.J. Pharmacological characterization of glycine-activated currents in HEK 293 cells expressing N-methyl-D-aspartate NR1 and NR3 subunits. J Pharmacol Exp Ther 322, 739–48 (2007).

9. Awobuluyi, M. et al. Subunit-specific roles of glycine-binding domains in activation of NR1/NR3 N-methyl-D-aspartate receptors. Mol Pharmacol 71, 112–22 (2007).

10. Kvist, T. et al. Crystal structure and pharmacological characterization of a novel N-methyl-D-aspartate (NMDA) receptor antagonist at the GluN1 glycine binding site. J Biol Chem 288, 33124–35 (2013).

11. Madry, C. et al. Principal role of NR3 subunits in NR1/NR3 excitatory glycine receptor function. Biochem Biophys Res Commun 354, 102–8 (2007).

12. Grand, T., Abi Gerges, S., David, M., Diana, M.A. & Paoletti, P. Unmasking GluN1/GluN3A excitatory glycine NMDA receptors. Nat Commun 9, 4769 (2018).

13. Rouzbeh, N. et al. Allosteric modulation of GluN1/GluN3 NMDA receptors by GluN1-selective competitive antagonists. J Gen Physiol 155(2023).

14. Wong, H.K. et al. Temporal and regional expression of NMDA receptor subunit NR3A in the mammalian brain. J Comp Neurol 450, 303–17 (2002).

15. Perez-Otano, I., Larsen, R.S. & Wesseling, J.F. Emerging roles of GluN3-containing NMDA receptors in the CNS. Nat Rev Neurosci 17, 623–35 (2016).

16. Das, S. et al. Increased NMDA current and spine density in mice lacking the NMDA receptor subunit NR3A. Nature 393, 377–81 (1998).

17. Roberts, A.C. et al. Downregulation of NR3A-containing NMDARs is required for synapse maturation and memory consolidation. Neuron 63, 342–56 (2009).

18. Murillo, A. et al. Temporal Dynamics and Neuronal Specificity of Grin3a Expression in the Mouse Forebrain. Cereb Cortex 31, 1914–1926 (2021).

19. Bossi, S. et al. GluN3A excitatory glycine receptors control adult cortical and amygdalar circuits. Neuron 110, 2438–2454 e8 (2022).

20. Yuan, T. et al. Expression of cocaine-evoked synaptic plasticity by GluN3A-containing NMDA receptors. Neuron 80, 1025–38 (2013).

21. Mueller, H.T. & Meador-Woodruff, J.H. NR3A NMDA receptor subunit mRNA expression in schizophrenia, depression and bipolar disorder. Schizophr Res 71, 361–70 (2004).

22. Lee, J.H., Zhang, J.Y., Wei, Z.Z. & Yu, S.P. Impaired social behaviors and minimized oxytocin signaling of the adult mice deficient in the N-methyl-d-aspartate receptor GluN3A subunit. Exp Neurol 305, 1–12 (2018).

23. Perez-Otano, I. et al. Assembly with the NR1 subunit is required for surface expression of NR3A-containing NMDA receptors. J Neurosci 21, 1228–37 (2001).

24. Al-Hallaq, R.A. et al. Association of NR3A with the N-methyl-D-aspartate receptor NR1 and NR2 subunits. Mol Pharmacol 62, 1119–27 (2002).

25. Zhu, Z. et al. Negative allosteric modulation of GluN1/GluN3 NMDA receptors. Neuropharmacology 176, 108117 (2020).

26. Otsu, Y. et al. Control of aversion by glycine-gated GluN1/GluN3A NMDA receptors in the adult medial habenula. Science 366, 250–254 (2019).

27. Bossi, S., Pizzamiglio, L. & Paoletti, P. Excitatory GluN1/GluN3A glycine receptors (eGlyRs) in brain signaling. Trends Neurosci 46, 667–681 (2023).

28. Lee, C.H. et al. NMDA receptor structures reveal subunit arrangement and pore architecture. Nature 511, 191–7 (2014).

29. Zhu, S. et al. Mechanism of NMDA Receptor Inhibition and Activation. Cell 165, 704–14 (2016).

30. Karakas, E. & Furukawa, H. Crystal structure of a heterotetrameric NMDA receptor ion channel. Science 344, 992–7 (2014).

31. Lu, W., Du, J., Goehring, A. & Gouaux, E. Cryo-EM structures of the triheteromeric NMDA receptor and its allosteric modulation. Science 355(2017).

32. Song, X. et al. Mechanism of NMDA receptor channel block by MK-801 and memantine. Nature 556, 515–519 (2018).

33. Jalali-Yazdi, F., Chowdhury, S., Yoshioka, C. & Gouaux, E. Mechanisms for Zinc and Proton Inhibition of the GluN1/GluN2A NMDA Receptor. Cell 175, 1520–1532 e15 (2018).

34. Chou, T.H., Tajima, N., Romero-Hernandez, A. & Furukawa, H. Structural Basis of Functional Transitions in Mammalian NMDA Receptors. Cell 182, 357–371 e13 (2020).

35. Wang, H. et al. Gating mechanism and a modulatory niche of human GluN1-GluN2A NMDA receptors. Neuron 109, 2443–2456 e5 (2021).

36. Chou, T.H., Kang, H., Simorowski, N., Traynelis, S.F. & Furukawa, H. Structural insights into assembly and function of GluN1-2C, GluN1-2A-2C, and GluN1-2D NMDARs. Mol Cell 82, 4548–4563 e4 (2022).

37. Zhang, J. et al. Distinct structure and gating mechanism in diverse NMDA receptors with GluN2C and GluN2D subunits. Nat Struct Mol Biol 30, 629–639 (2023).

38. Meyerson, J.R. et al. Structural mechanism of glutamate receptor activation and desensitization. Nature 514, 328–34 (2014).

39. Meyerson, J.R. et al. Structural basis of kainate subtype glutamate receptor desensitization. Nature 537, 567–571 (2016).

40. Khanra, N., Brown, P.M., Perozzo, A.M., Bowie, D. & Meyerson, J.R. Architecture and structural dynamics of the heteromeric GluK2/K5 kainate receptor. Elife 10(2021).

41. Michalski, K. & Furukawa, H. Structure and function of GluN1-3A NMDA receptor excitatory glycine receptor channel. Sci Adv 10, eadl5952 (2024).

42. Yao, Y. & Mayer, M.L. Characterization of a soluble ligand binding domain of the NMDA receptor regulatory subunit NR3A. J Neurosci 26, 4559–66 (2006).

43. Sheldon, N. et al. Generation of Conformation-Specific Monoclonal Antibodies for Integral Membrane Proteins. Curr Protoc 5, e70142 (2025).

44. Hattori, M., Hibbs, R.E. & Gouaux, E. A fluorescence-detection size-exclusion chromatography-based thermostability assay for membrane protein precrystallization screening. Structure 20, 1293–9 (2012).

45. Jain, A. et al. Probing cellular protein complexes using single-molecule pull-down. Nature 473, 484–8 (2011).

46. Wang, T.M. et al. A novel NMDA receptor positive allosteric modulator that acts via the transmembrane domain. Neuropharmacology 121, 204–218 (2017).

47. Atomwise, A.P. AI is a viable alternative to high throughput screening: a 318-target study. Sci Rep 14, 7526 (2024).

48. Furukawa, H., Singh, S.K., Mancusso, R. & Gouaux, E. Subunit arrangement and function in NMDA receptors. Nature 438, 185–92 (2005).

49. Gielen, M., Siegler Retchless, B., Mony, L., Johnson, J.W. & Paoletti, P. Mechanism of differential control of NMDA receptor activity by NR2 subunits. Nature 459, 703–7 (2009).

50. Yao, Y., Harrison, C.B., Freddolino, P.L., Schulten, K. & Mayer, M.L. Molecular mechanism of ligand recognition by NR3 subtype glutamate receptors. EMBO J 27, 2158–70 (2008).

51. Yao, Y., Belcher, J., Berger, A.J., Mayer, M.L. & Lau, A.Y. Conformational analysis of NMDA receptor GluN1, GluN2, and GluN3 ligand-binding domains reveals subtype-specific characteristics. Structure 21, 1788–99 (2013).

52. Chou, T.H. et al. Molecular mechanism of ligand gating and opening of NMDA receptor. Nature 632, 209–217 (2024).

53. Twomey, E.C., Yelshanskaya, M.V., Grassucci, R.A., Frank, J. & Sobolevsky, A.I. Channel opening and gating mechanism in AMPA-subtype glutamate receptors. Nature 549, 60–65 (2017).

54. Wada, A., Takahashi, H., Lipton, S.A. & Chen, H.S. NR3A modulates the outer vestibule of the “NMDA” receptor channel. J Neurosci 26, 13156–66 (2006).

55. Kang, H. et al. Mechanism of conductance control and neurosteroid binding in NMDA receptors. Nature 648, 220–228 (2025).

56. Abbott, J.A. et al. Cryo-EM snapshots of NMDA receptor activation illuminate sequential rearrangements. Sci Adv 11, eadx4647 (2025).

57. Amin, J.B., Gochman, A., He, M., Certain, N. & Wollmuth, L.P. NMDA Receptors Require Multiple Pre-opening Gating Steps for Efficient Synaptic Activity. Neuron 109, 488–501 e4 (2021).

58. Twomey, E.C., Yelshanskaya, M.V., Grassucci, R.A., Frank, J. & Sobolevsky, A.I. Structural Bases of Desensitization in AMPA Receptor-Auxiliary Subunit Complexes. Neuron 94, 569–580 e5 (2017).

59. Gangwar, S.P. et al. Kainate receptor channel opening and gating mechanism. Nature 630, 762–768 (2024).

60. Pintilie, G. et al. Measurement of atom resolvability in cryo-EM maps with Q-scores. Nat Methods 17, 328–334 (2020).

## References

61. Goehring, A. et al. Screening and large-scale expression of membrane proteins in mammalian cells for structural studies. Nat Protoc 9, 2574–85 (2014).

62. Punjani, A., Zhang, H. & Fleet, D.J. Non-uniform refinement: adaptive regularization improves single-particle cryo-EM reconstruction. Nat Methods 17, 1214–1221 (2020).

63. Punjani, A., Rubinstein, J.L., Fleet, D.J. & Brubaker, M.A. cryoSPARC: algorithms for rapid unsupervised cryo-EM structure determination. Nat Methods 14, 290–296 (2017).

64. Zivanov, J. et al. New tools for automated high-resolution cryo-EM structure determination in RELION-3. Elife 7(2018).

65. Pettersen, E.F. et al. UCSF Chimera--a visualization system for exploratory research and analysis. J Comput Chem 25, 1605–12 (2004).

66. Jumper, J. et al. Highly accurate protein structure prediction with AlphaFold. Nature 596, 583–589 (2021).

67. Varadi, M. et al. AlphaFold Protein Structure Database: massively expanding the structural coverage of protein-sequence space with high-accuracy models. Nucleic Acids Res 50, D439–D444 (2022).

68. Liebschner, D. et al. Macromolecular structure determination using X-rays, neutrons and electrons: recent developments in Phenix. Acta Crystallogr D Struct Biol 75, 861–877 (2019).

69. Emsley, P., Lohkamp, B., Scott, W.G. & Cowtan, K. Features and development of Coot. Acta Crystallogr D Biol Crystallogr 66, 486–501 (2010).

70. Krissinel, E. & Henrick, K. Inference of macromolecular assemblies from crystalline state. J Mol Biol 372, 774–97 (2007).

71. Pettersen, E.F. et al. UCSF ChimeraX: Structure visualization for researchers, educators, and developers. Protein Sci 30, 70–82 (2021).

72. Zhong, E.D., Bepler, T., Berger, B. & Davis, J.H. CryoDRGN: reconstruction of heterogeneous cryo-EM structures using neural networks. Nat Methods 18, 176–185 (2021).

73. Clark, S. et al. Single molecule studies of the native hair cell mechanosensory transduction complex. bioRxiv (2023).

74. Chen, C. & Okayama, H. High-efficiency transformation of mammalian cells by plasmid DNA. Mol Cell Biol 7, 2745–52 (1987).

75. Christopoulos, A. Assessing the distribution of parameters in models of ligand-receptor interaction: to log or not to log. Trends Pharmacol Sci 19, 351–7 (1998).

